# This must be the place: deep learning local adaptation

**DOI:** 10.64898/2026.09.16.752190

**Authors:** Jordan Rodriguez, Richard Cronn, Silas Tittes, Andrew D. Kern

## Abstract

Climate change is increasingly disrupting the relationship between locally adapted populations and the environments in which they evolved, creating an urgent need for tools that connect genomic variation to climate. Common-garden and provenance trials remain the gold standard for characterizing local adaptation, but their time and resource requirements limit how broadly they can be applied. Genomic approaches provide a complementary path. Genotype–environment association (GEA) methods identify environmentally associated loci. Machine-learning models have also shown that geographic origin can be predicted directly from genotypes. Here we introduce `EcoLocator`, a supervised deep neural network that jointly predicts geographic location and climate of origin from genotypes. Through extensive simulations we demonstrate that `EcoLocator` accurately recovers geographic location and environment of origin from genotype data, and, with SHAP-based feature attribution, identifies adaptive loci more reliably than benchmark GEA methods. We apply our method to coastal Douglas-fir (*Pseudotsuga menziesii* var. *menziesii*), where `EcoLocator` predicts geographic origin (*R*^2^ = 0.75–0.83) and climate of origin (*R*^2^ = 0.52–0.72) under leave-one-out cross-validation. Notably, we find that climate predicted directly from genotypes outperforms climate inferred by first predicting geographic origin, showing that `EcoLocator` captures genotype–climate signal that cannot be recovered from geography alone. Our prediction errors fall within the tolerances used in existing seed-transfer guidelines, demonstrating that `EcoLocator`’s predictions are ready for practical application, and our approach is readily extendable to other species and conservation contexts.

## Introduction

Population genetic theory predicts that the distribution of genetic variation within and between populations is shaped by demographic history, spatial structure, and selective pressures, and thus carries information about the processes that generate and maintain it (Lewontin, 1974; Hartl and Clark, 1980; Ellegren and Galtier, 2016). In spatially structured, locally adapted populations, this variation encodes information about both geographic location and climate of origin. Geographic structure arises because local mating and limited dispersal generate isolation by distance, causing alleles to be more similar among neighboring individuals than among those farther apart (Wright, 1943; Slatkin, 1993). Environmental selection, in contrast, produces allele frequency clines that parallel ecological gradients, as individuals carrying locally beneficial variants display higher fitness in particular environments (Endler, 1977; Clausen, Keck, and Hiesey, 1948; Kawecki and Ebert, 2004). In principle the two signals can be told apart, since neutral structure follows geographic distance while adaptive signals follow particular environmental variables such as temperature, moisture, or seasonality (Rellstab et al., 2015; Hoban et al., 2016; Tigano and Friesen, 2016; Coop et al., 2010). In practice, the separation is never clean. Range expansions, bottlenecks, and drift can all produce allele frequency gradients that look like adaptive clines (Excoffier, Foll, and Petit, 2009; Novembre and Di Rienzo, 2009), while spatially autocorrelated environmental variables can make real adaptive gradients look like isolation by distance, inflating false-positive rates in tests used to distinguish them (e.g., Guillot and Rousset, 2013; Legendre, Fortin, and Borcard, 2015). Here we ask whether a single predictive model, trained directly on genomic data, can learn to tell these two signals apart on its own.

Local adaptation, in which populations evolve higher fitness in their local environments, can generate covariation of allele frequencies with environmental gradients when spatially heterogeneous selection outweighs the homogenizing effects of gene flow (Savolainen, Lascoux, and Merilä, 2013; Lenormand, 2002). Beyond increasing local fitness, local adaptation sustains genetic diversity across heterogeneous landscapes by maintaining quantitative genetic variance and, at some loci, multiple adaptive alleles, helping populations retain the variation on which selection can act as environments change (Hereford, 2009). The classic models of Wright (1943), Levene (1953), and Endler (1977) work out how selection, migration, and drift combine to produce predictable spatial patterns of variation, and later work has filled in the details. Even modest selection can create steep genomic clines when dispersal is limited (Tigano and Friesen, 2016; Yeaman and Whitlock, 2011), and strong gene flow will swamp adaptive alleles unless they are tightly linked or of large effect (Slatkin, 1975). Climate change is now pulling populations out of step with the environments they are adapted to (Calvin et al., 2023), and as ranges shift, managers and stakeholders need to know where populations will persist under conditions they have not experienced before. Most of what we have learned about local adaptation comes from common-garden experiments, reciprocal transplants, and provenance tests; however these approaches are slow and expensive, and for most species they have never been done (Aitken and Whitlock, 2013). Genomic data are cheaper and faster to collect, and the field of landscape genomics has grown up around the effort to connect spatial and environmental variation with patterns of genetic diversity (Manel et al., 2003; Rellstab et al., 2015). In forest trees these methods are already starting to inform seed transfer and assisted gene flow decisions (Aitken and Bemmels, 2016).

Most landscape genetics work relies on genotype–environment association (GEA) methods, which identify putatively adaptive loci by testing for statistical associations between allele frequencies and environmental variables (Rellstab et al., 2015). Methods such as `Bayenv2` (Günther and Coop, 2013), latent factor mixed models (Frichot et al., 2013), and `BayPass` (Gautier, 2015) test loci one at a time against environmental predictors, typically while controlling for population structure through covariance matrices or latent factors. While widely applied, these methods are limited in their ability to capture polygenic adaptation involving many loci of small effect, and the neutral gradients described above can produce false positives (Lotterhos and Whitlock, 2015). Newer approaches address some of these limitations by analysing many loci and environmental predictors simultaneously. Redundancy analysis (RDA) has proven effective at detecting multilocus adaptive signatures (Forester et al., 2018; Capblancq and Forester, 2021), and gradient forest methods model nonlinear allele–environment relationships across the genome (Fitzpatrick and Keller, 2015). More recently, the Weighted-Z Analysis (`WZA`) introduced a window-based framework for aggregating single-locus GEA statistics into region-level summaries of adaptive signal (Booker et al., 2024). All of these methods, old and new, are built to *test* for associations between genotype and environment. Spatial population structure enters as a nuisance to be corrected for, not as something worth modeling in its own right. So none of them will tell you where an individual came from, and with the partial exception of gradient-forest and genomic-offset approaches, which project allele turnover under new climates (Fitzpatrick and Keller, 2015), none of them will predict an individual’s environment of origin from its genotype either.

Machine learning (ML) offers a different way in. Rather than testing loci one at a time, a supervised model can use the whole genotype at once to predict geographic origin or climate of origin directly. ML methods can also be effective at pulling patterns out of high-dimensional data with nonlinear interactions among loci and correlated predictors (Schrider and Kern, 2018), though performance depends on how well the training data represent the populations and conditions where the model is applied. Battey, Ralph, and Kern (2020a) demonstrated that a deep neural network, `Locator`, can predict the geographic origin of an individual from its genotype alone by learning from multilocus patterns of isolation by distance. `Locator` achieves high spatial accuracy, but it does not model environmental covariates, so its predictions give a climate of origin only indirectly, through a lookup at the predicted location. While a location prediction implies a climate of origin once paired with a map of historical climate normals, that inference assumes the climate at a given location stays put — an assumption that erodes as climates shift faster than the normals they’re being compared against. Moreover, these prediction offer nothing where future conditions have no analog anywhere on today’s landscape. Predicting climate of origin directly from genotype, rather than through an intermediate geographic prediction, sidesteps that assumption.

Other recent work brings in ecology more directly. Gradient forest methods link genomic variation to environmental predictors to forecast climate-driven maladaptation (Fitzpatrick, Chhatre, et al., 2021). In crop genomics, for example, deep learning models that take in weather, soil, and other covariates improve predictions of how a genotype will perform across climates (Washburn et al., 2021; Montesinos-Ĺopez et al., 2021). No existing method, though, predicts geography and environment from genotype together, in one model that can learn from neutral spatial structure and adaptive environmental associations at the same time.

Here we introduce `EcoLocator`, a deep neural network that extends Locator by jointly predicting geographic location and climate of origin from genomic data. Trained on georeferenced genotypes paired with historical climate data, the model learns multilocus associations with both geographic position and environmental gradients, so that it can infer climate of origin directly from genotype when allele frequencies covary with climate. We pair the model with a feature attribution method, SHapley Additive exPlanations (SHAP) (Lundberg and S.-I. Lee, 2017), to see which input SNPs each prediction depends on, and so which variants contribute most to adaptive differentiation across the landscape. We test `EcoLocator` on simulations to find out where it works well and where it does not, then apply it to an empirical dataset of coastal Douglas-fir (*Pseudotsuga menziesii* var. *menziesii*).

Coastal Douglas-fir is a member of the pine family with a lineage dating to the Paleocene (Hermann, 1985) and a natural range spanning extensive climatic gradients along the North American Pacific coast, from California to British Columbia. Although most populations are characterized by low genetic differentiation, high gene flow, and high outcrossing rates (Slavov, Howe, and Adams, 2005), they still exhibit strong signatures of local adaptation, with adaptive quantitative phenotypes tracking climatic features such as winter cold temperatures and summer aridity (St. Clair, Mandel, and Vance-Borland, 2005). A century of provenance trials and seed-transfer research has shown that trees express maladaptive responses (e.g., slow growth, poor form, disease susceptibility) when moved beyond their local source climate, with home-environment advantage and genotype–environment interactions structured primarily by winter cold, summer drought, and summer heat accumulation (St. Clair, Howe, and Kling, 2020; St. Clair, Mandel, and Vance-Borland, 2005; St. Clair and Howe, 2007). To minimize maladaptation, seed zones define the geographic limits within which seedlings can be moved while preserving local adaptation. Douglas-fir seed zones were formalized for U.S. Forest Service lands in the 1940s (Kummel, Rindt, and Munger, 1944), expanded to federal, state, and private lands in the 1960s and 1970s (USDA Forest Service, 1973), and later revised to incorporate climatic factors and experimental evidence from reciprocal transplant trials (e.g., Randall, 1996; St. Clair, Richardson, et al., 2022). Seed zones also give us something simulations cannot, which is a standard already in use by managers for deciding whether a predicted climate is close enough to the true one. This combination of strong local adaptation and a long history of ecological and silvicultural study makes Douglas-fir a natural bridge between controlled simulations and real-world management applications.

To our knowledge `EcoLocator` is the first open-source tool that infers both the geographic origin of a genotype and the climate it is adapted to, which is the information managers will need as climates shift faster than provenance trials can keep up.

### EcoLocator

`EcoLocator` is an open-source Python package for predicting the geographic origin and climate of origin of individual genotypes, accessible via both a command-line interface and a programmatic Python API. ^1^ `EcoLocator` does not depend on fixed assumptions about allele–environment relationships, but learns directly from high-dimensional genomic and environmental data (Figure 1). By jointly predicting location and environmental variables, it captures both spatial genetic structure and signals of local adaptation, providing a scalable, biologically informed approach for studying complex evolutionary patterns. Both the CLI and Python API accept standard genotype formats (VCF, Zarr, tab-delimited matrices) as input, facilitating integration into existing genomic workflows whether scripting interactively or from the command line.

**Figure 1:**
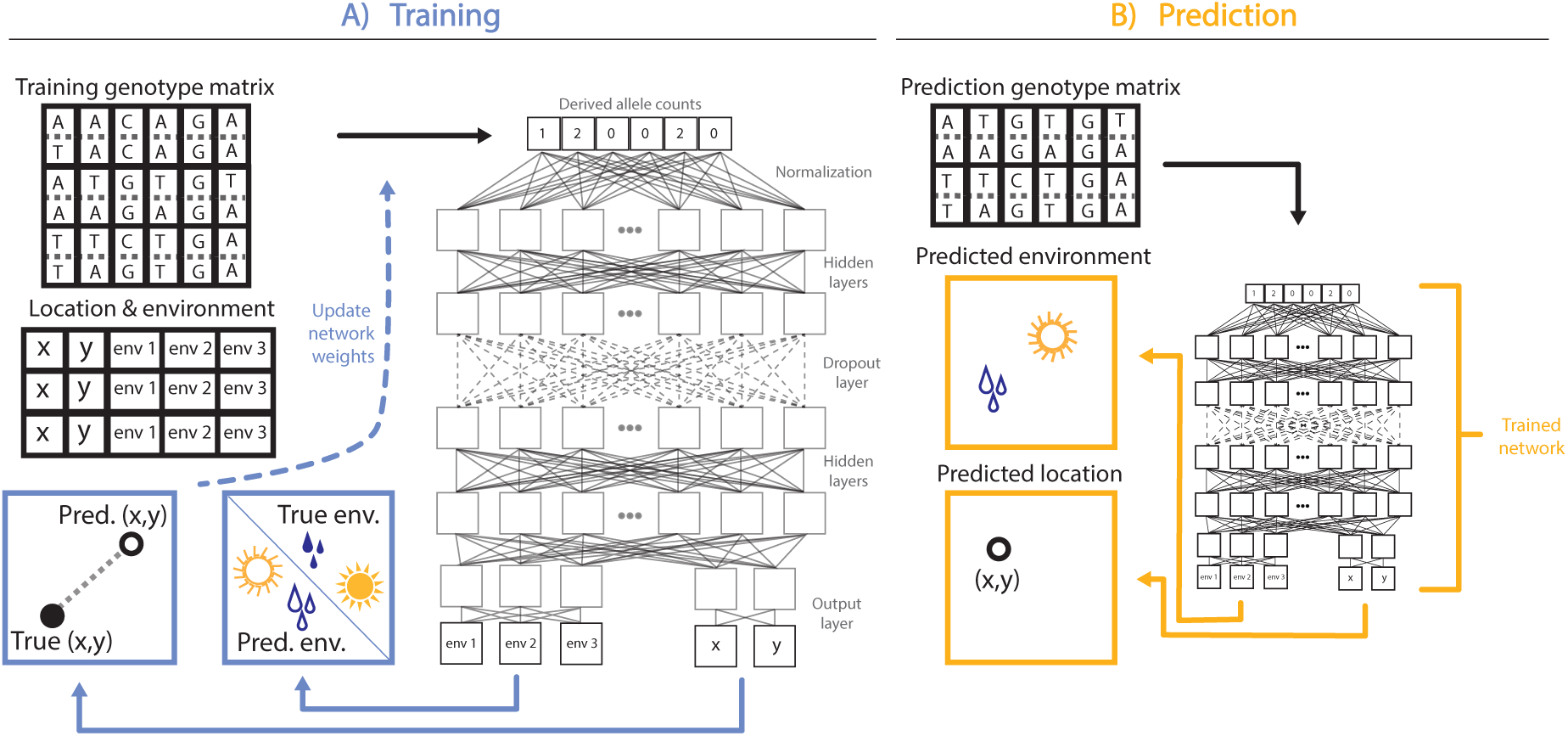
`EcoLocator` architecture. **Panel A** shows the training regime, with the inputs traversing the network in a supervised fashion; network weights are updated through backpropagation until predictions are optimized toward the true value, with the process shown here in blue. Once trained, `EcoLocator` enters the prediction step, shown here in **Panel B**. Given genotypes alone, the trained network jointly predicts climate of origin and geographic location, shown in yellow.

### Preprocessing

`EcoLocator` converts genomic data from VCF, Zarr, and tab-delimited matrix formats into vectors of minor allele counts per individual using the scikit-allel (Miles et al., 2024) and NumPy (Oliphant and Team, 2024) libraries. Missing genotypes are imputed by drawing alleles from a binomial distribution parameterized by the site’s allele frequency, an approximation that preserves expected variation while minimizing bias. Sites can be filtered by a minor allele count threshold to reduce noise from rare variants that contribute little predictive signal. Geographic coordinates are normalized to zero mean and unit variance, and environmental covariates are standardized similarly prior to training.

### Model architecture

`EcoLocator` employs a multilayer perceptron with a shared trunk and two task-specific output heads, trained under a supervised learning framework on labeled datasets of genotypes paired with known geographic coordinates and climate variables. Input genotypes pass through a batch normalization layer before entering the shared trunk, which consists of dense layers with ELU activations and a central dropout layer for regularization. The trunk branches into two heads — one predicting 2D geographic coordinates (i.e., latitude and longitude), one predicting environmental covariates — each implemented as a small two-layer subnetwork. The network is optimized using the Adam optimizer with a custom multitask loss function, combining mean Euclidean distance on the standardized latitude and longitude for geographic prediction and MSE on the standardized environmental covariates, each scaled by an independently specified weight. Concretely, the loss function is:

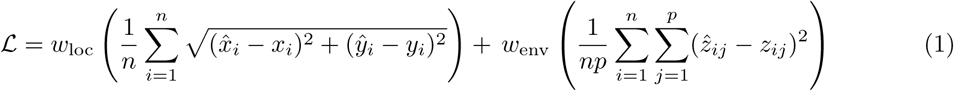

where *n* is the number of training samples, *p* is the number of environmental covariates, *x_i_* and *y_i_* are the true longitude and latitude of sample *i*, *x̂_i_* and *ŷ_i_* are the predicted coordinates, *z_ij_* is the true value of covariate *j* for sample *i*, *ẑ_ij_* is the predicted value, and *w*_loc_*, w*_env_ *≥* 0 are loss weights (both defaulting to 1.0, see Figure S1). The network is trained using a standard validation scheme in which a user-specified fraction of samples (default 0.9) is used for model fitting, while the remaining fraction is held out for validation, enabling convergence monitoring, early stopping, and adaptive learning rate reduction to prevent overfitting. By explicitly coupling geographic and climatic inference, the model captures both neutral spatial structure and environment-associated genomic variation.

In practice, an end-to-end `EcoLocator` analysis consists of five steps:

1. Assemble a training dataset consisting of filtered genotype data in a supported format (VCF, Zarr, or tab-delimited matrix) and a sample metadata file pairing each individual with its known geographic coordinates and environmental covariate values. Samples to be predicted should have their coordinates and covariates left blank.
2. Train the model using `EcoLocator` train (or the Python API), specifying the genotype file, sample metadata, and an output directory. The model is fit to jointly predict geographic origin and environmental covariates, with early stopping and adaptive learning rate reduction applied automatically.
3. Evaluate model performance using `EcoLocator` loo, which iteratively holds out each sample with known coordinates, trains on the remainder, and predicts the held-out individual (leave-one-out cross-validation). The resulting predictions can be compared against true values to assess spatial and environmental accuracy before applying the model to unknown samples.
4. Predict geographic origin and environmental covariates for samples with unknown locations using `ecolocator predict`, providing the saved model directory and a genotype file containing the target samples.
5. (**Optional**) Quantify per-SNP importance for predicted samples using `EcoLocator` attribute, which applies SHAP values to identify loci driving each prediction.

The predicted coordinates and environmental values from steps 2–4 provide continuous estimates of geographic and climatic origin, and SHAP attributions from step 5 can be used to identify loci contributing most strongly to each prediction. More details are given below.

## Methods

### Simulating local adaptation in continuous space (non-Wright–Fisher)

To explore under which evolutionary scenarios `EcoLocator` performs best, we used `SLiM v.4.2.2` (Haller and Messer, 2023) to simulate a spatially explicit non-Wright–Fisher population experiencing local adaptation of a quantitative trait that evolves to track a single continuously varying environmental axis. By varying simulation parameters such as dispersal, genetic architecture of the quantitative trait, and the strength of stabilizing selection, the evolutionary trajectory of a given population can be changed in predictable ways. For example, when environmental selection is weak and individuals disperse far distances, the homogenizing effects of gene flow across the landscape result in little correlation between genetic variation and environmental variation (Lenormand, 2002; Kawecki and Ebert, 2004; Booker et al., 2024). The underlying spatial and density-dependent components of the simulation were borrowed from Chevy et al. (2025), but are briefly described below for completeness.

### Fitness and reproduction

To capture the joint effects of selection, dispersal, and spatial dynamics, we simulate individuals in continuous space on top of an environmental grid (see Figure S7). We initialize each simulation by randomly placing individuals across the landscape. Individuals occupy a continuous, square landscape, which we divide into an 8×8 grid of equal-sized regions, each assigned its own environmental value that defines the local phenotypic optimum, *θ*, for individuals within it. Values are generated using the midpoint-displacement neutral landscape model implemented in the NLMpy Python package (Etherington, Holland, and O’Sullivan, 2015), with a Hurst exponent of zero (H=0), which removes spatial autocorrelation and creates a patchwork of optima. This setup allows us to isolate the effects of spatial dynamics and dispersal from those of environment-dependent selection. Critically, because grid cells have environmental optima that are uncorrelated at the scale of the grid, proximity on the map does not imply similarity in environment: individuals on opposite sides of the landscape can share the same optimum, while individuals in adjacent cells can be adapted to entirely different ones. Geographic and environmental prediction are therefore decoupled; therefore, any accuracy `EcoLocator` achieves on the environmental target cannot arise from geographic proximity or from neutral spatial structure driven by isolation by distance, but must instead reflect genotype–environment covariance generated by local adaptation.

Individuals experience fitness as a combination of stabilizing selection and density-dependent competition. Stabilizing selection follows a Gaussian fitness function centered on the local optimum (Equation (2)), where *z* is the individual’s phenotype, *θ* is the optimum in the corresponding grid cell, and *σ_K_* controls the strength of selection:

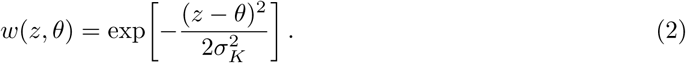

Total fitness is then determined by multiplying this term with a density-dependent factor *C_i_*, where competition is implemented as a Beverton–Holt style density regulation function,

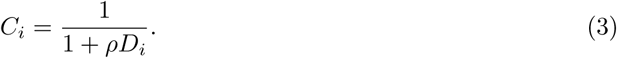

where *D_i_* is the local density of neighbors around individual *i*, computed as a Gaussian-weighted count of nearby individuals, and *ρ* scales the strength of competition. Individuals in crowded regions are therefore less likely to survive and reproduce.

After survival, individuals disperse according to a dispersal kernel that sets the spatial scale of gene flow, redistributing individuals across the landscape. Following dispersal, individuals mate with nearby partners, so that reproduction depends on both spatial proximity and local density. By varying the dispersal parameter and the standard deviation of the fitness function, we can impact the strength of local adaptation in the simulation (Figure 2, Figure S4). Together, stabilizing selection, competition, dispersal, and local mating govern the survival and reproduction of individuals.

**Figure 2:**
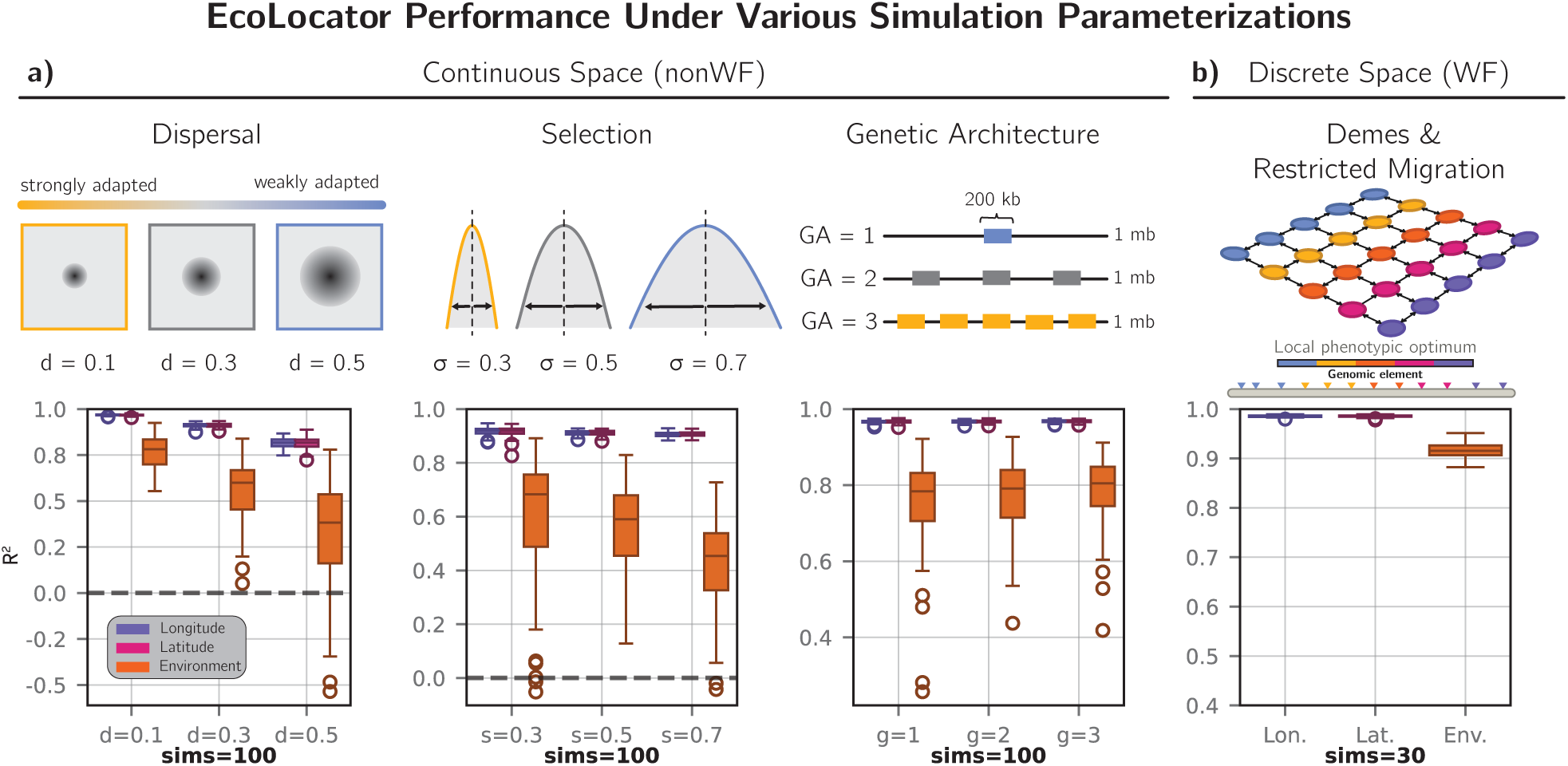
Effects of parameter variation on `EcoLocator` performance. **Panel a)** shows the continuous-space, non-Wright–Fisher simulation parameterizations in the top row, with prediction accuracy for each respective parameter shown as boxplots in the bottom row. We varied dispersal (*σ_D_* = 0.1, 0.3, 0.5), selection strength (*σ_K_* = 0.3, 0.5, 0.7), and genetic architecture (GA = 1, 2, 3). Yellow indicates parameter regimes expected to produce stronger local adaptation, including limited dispersal and stronger selection, whereas blue indicates parameter regimes expected to produce weaker local adaptation, including higher dispersal and weaker selection. Boxplots show prediction accuracy, measured as *R*^2^ using scikit-learn, for longitude, latitude, and environmental targets across the varied simulation parameters. Each continuous-space simulation parameter regime was replicated 100 times. **Panel b)** shows the discrete-space, Wright–Fisher simulation regime in the top row, with corresponding prediction accuracy shown in the bottom row. The Wright–Fisher simulations were held under a fixed parameterization and replicated 30 times, providing a complementary comparison to the continuous-space parameter sweeps. Boxplots summarize `EcoLocator` prediction accuracy across geographic and environmental targets in this discrete-deme framework.

Local adaptation can then be quantified as the mean absolute difference between each individual’s phenotype and its local optimum, such that smaller values correspond to stronger local adaptation:

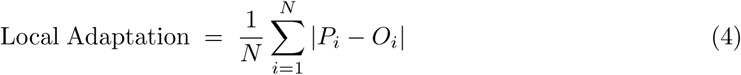

where *P_i_* is the phenotype of individual *i*, *O_i_* is the local optimum at individual *i*’s location, and *N* is the total number of individuals. Because dispersal, selection strength, and genetic architecture interact to determine how much local adaptation actually emerges, this metric lets us measure the realized outcome of each parameter combination directly, rather than inferring it from the parameters alone, so that it can be related to `EcoLocator`’s prediction accuracy.

### Genomic architecture and tree sequence recording

The genetics underlying adaptation are simulated using a 1 Mb chromosome carried by each individual, along which mutations at quantitative trait loci (QTL) arise, with additive effects that determine an individual’s phenotype. Because the simulated chromosome spans only 1 Mb, a per-base recombination rate of 10*^−^*^8^ produces very little recombination across it and consequently extensive linkage. We therefore used 10*^−^*^6^, which yields approximately one expected recombination breakpoint across the chromosome per meiosis, reducing long-range LD and allowing clearer evaluation of locus-level feature attribution (Figure S2).

To explore whether the genomic distribution of adaptive loci influences the detectability of local adaptation, we varied the arrangement of QTLs along the chromosome across three genetic architectures. In the first, all QTLs were clustered in a single 200 kb window (bases 400,000–599,999; spanning 20% of the chromosome), centered at the midpoint of the chromosome. In the second, QTLs were split across three evenly-spaced 100 kb windows (10% of the chromosome each, 30% total span), centered at bases 250,000, 500,000, and 750,000. In the third, QTLs were split across five evenly-spaced 100 kb windows (10% of the chromosome each, 50% total span), centered at approximately bases 166,667, 333,333, 500,000, 666,667, and 833,333.

These treatments allowed us to test whether adaptive signals are easier to detect when concentrated in specific genomic regions versus dispersed more broadly (Figure 2, panel a). We only simulate non-neutral mutations within SLiM. Genomic information is stored in tree sequences and recapitated using pyslim 1.0.4 (Haller, Galloway, et al., 2019). Neutral mutations are then overlaid onto the recapitated tree sequences using msprime v.1.1.1 (Baumdicker et al., 2022), allowing us to track both adaptive and neutral background variation separately.

### Varying dispersal, selection, and genetic architecture parameters

To evaluate the robustness of `EcoLocator` across diverse evolutionary contexts, we varied three parameters: dispersal, the strength of stabilizing selection, and the genomic architecture of adaptive loci (described above). We picked these three because each one governs a process that sets how detectable local adaptation is. Gene flow homogenizes allele frequencies across space, selection pulls populations in different environments apart, and genetic architecture decides where in the genome the adaptive alleles sit. Dispersal distance (*σ_D_*) sets the spatial scale of gene flow. Small values keep offspring near their parents and let neighboring regions differentiate; large values mix individuals across the landscape. We tested *σ_D_* = 0.1, 0.3, 0.5, which for reference correspond to 2%, 6%, and 10% of the landscape width moved per generation, since the landscape spans 5 *×* 5 spatial units. The strength of stabilizing selection is set by the standard deviation (*σ_K_*) of the Gaussian fitness function. Small *σ_K_* punishes phenotypes far from the local optimum, while large *σ_K_* tolerates them. We tested *σ_K_* = 0.3, 0.5, 0.7 for strong, intermediate, and weak selection. When varying one parameter we held the others fixed: the dispersal sweep used *σ_K_* = 0.5 and genetic architecture 1, the selection sweep used *σ_D_* = 0.3 and genetic architecture 1, and the architecture sweep used *σ_D_* = 0.1 and *σ_K_* = 0.5. Spatial and phenotypic scales in these simulations are defined relative to the simulated landscape and environmental gradient rather than to physical distances, and are therefore best interpreted as ratios. Mutation and recombination rates, by contrast, are in standard per-base, per-generation units. Taken together these runs span populations that are highly structured and strongly adapted at one end, and weakly structured and weakly adapted at the other, which is the range over which we wanted to see where `EcoLocator` works and where it does not.

### Simulating local adaptation in discrete space (Wright–Fisher)

To complement our spatially explicit simulations, we implemented a second set of forward-in-time simulations based on the Wright–Fisher framework described in Booker et al. (2024). Simulation design and parameterization closely followed the implementation provided in the `WZA` GitHub repository (https://github.com/TBooker/WZA/tree/master), with minor modifications to ensure compatibility with the current version of `F`. These simulations were designed to more closely mirror the evolutionary scenarios used to evaluate `F`, allowing for a more direct comparison between `F`and existing GEA methods. Following Booker et al. (2024), we simulated a two-dimensional stepping-stone metapopulation composed of discrete demes arranged on a grid, with a constant population size per deme (*N* = 100) under Wright–Fisher dynamics. Migration was restricted to neighboring demes at a rate of *m* = 0.0375 per generation in each cardinal direction (demes at grid edges and corners had correspondingly fewer migrant sources), producing a pattern of isolation by distance and spatial genetic structure across the landscape. Environmental heterogeneity was defined using the British Columbia (BC) map of climatic variation, which assigns each deme a local phenotypic optimum based on discretized environmental values. In contrast to our main simulations, where a continuous phenotype is pulled toward a local optimum that varies across the landscape and fitness declines with deviation from that optimum in either direction, these Wright–Fisher simulations implement directional selection at a small number of discrete loci: at each locus, one allele is simply favored over the other depending on the local environment, and fitness increases monotonically with the number of copies carried, with no penalty for carrying “too many.” Specifically, 12 biallelic loci distributed across the genome experience genic selection, in which each copy of the derived allele multiplies fitness by (1 + *sθ*), where *θ* is the local phenotypic optimum assigned to that deme and *s* is a fixed selection coefficient (*s* = 0.0136), with heterozygotes receiving half this effect. Because *θ* is positive in demes where the derived allele is advantageous and negative where it is disadvantageous, selection at each locus consistently favors whichever allele matches the local environment. This formulation produces strong genotype–environment associations localized to a relatively small number of genomic regions.

Simulations were run in two phases: an initial neutral burn-in period to establish population structure and linkage patterns, followed by a directional selection phase during which adaptive alleles increase in frequency in a spatially heterogeneous manner. Tree-sequence recording was used to capture the full genealogical history of each simulation, and neutral mutations were subsequently overlaid to generate genome-wide variation. To prepare as input to `F`, deme coordinates were encoded as latitude and longitude, using the row and column indices of each deme in the grid, and the environmental variable was defined as the local phenotypic optimum for each deme. Genotype matrices were extracted from tree sequences and used as input for both `F`and downstream GEA analyses. With few loci under directional selection and structure imposed by discrete demes, these simulations are a simpler setting than our continuous-space ones, with genotype–environment associations that are strong and confined to a small part of the genome. They give us a useful contrast to the harder spatial scenarios.

### Feature attribution

To quantify the contribution of individual genetic features to `F`’s predictions, we use the `F`Python package (`f`*≥*`f`) (Lundberg and S.-I. Lee, 2017), which assigns importance scores based on each feature’s marginal contribution to the model output across all possible feature coalitions. Attributions were computed separately for each prediction target (latitude, longitude, and environmental covariates), allowing the genetic contributions to be evaluated conditional on the response variable of interest. SHAP values were calculated on trained models using the genotype matrix as input, attributing predictive signal to individual loci while accounting for nonlinear interactions among sites. To complement SHAP-based attribution and assess the robustness of inferred signals, we performed a series of permutation-based sensitivity analyses, testing whether `F`’s accuracy depends primarily on the selected QTL window, SNPs in linkage disequilibrium with that window, or signal distributed across the genotype matrix genome-wide (Figure S3). In the genotype matrix, with individuals as rows and SNPs as columns, we independently shuffled each column’s values across individuals prior to prediction, breaking that SNP’s association with each individual’s true label while leaving its allele frequency and every other column unchanged; because each column was permuted independently, this also destroyed any correlation structure among jointly permuted sites. These permutations included random shuffling of increasing proportions of sites genome-wide, targeted permutations of predefined genomic windows containing selected variants, and permutations of loci in linkage disequilibrium (LD) with those windows. LD partners were identified using pairwise correlation thresholds (see Figure S2), and permutations were applied either to the selected window alone or jointly with subsets of LD-linked sites stratified by LD strength. This framework enables systematic evaluation of how predictive performance responds to disruption of adaptive loci, correlated hitchhiking regions, and broader neutral genetic structure, providing an interpretable link between model predictions, genetic architecture, and simulated evolutionary processes. Feature attribution analyses were performed separately for each simulation regime, allowing us to assess how the recovery of adaptive loci depends on the underlying evolutionary scenario.

#### Empirical FDR scores from SHAP

SHAP values do not inherently provide a measure of statistical significance, as they represent model-derived feature importance scores rather than hypothesis test statistics. To enable direct comparison with GEA methods, which report *p*-values for association, we transformed SHAP-based rankings into an empirical FDR score framework. SHAP values were first aggregated across samples by computing the mean absolute SHAP value per SNP. Variants were then ranked from highest to lowest based on this aggregated score. To enable comparison with GEA-derived *p*-values, we calculated an empirical FDR score for each variant using a rank-based framework. At each rank position *i*, all variants up to that point were treated as significant, and the proportion of false positives (neutral variants) relative to the total number of variants in the set was calculated. This assigns each SNP an empirical FDR score based on its SHAP threshold. Because this approach relies on known neutrality labels, these FDR scores represent an empirical, supervised estimate of significance rather than a model-based or permutation-based test.

### Comparing EcoLocator to other GEA methods

To benchmark the ability of `EcoLocator`-derived feature attribution to identify adaptive loci, we compared SHAP-based variant rankings to two genome–environment association (GEA) approaches: the Weighted-Z Analysis (`WZA`) (Booker et al., 2024) and the top-candidate method (Yeaman, Hodgins, et al., 2016). These comparisons were performed across both simulation regimes described above, allowing us to evaluate method performance under distinct evolutionary scenarios. `WZA` and the top-candidate method are both window-based approaches designed to capture this regional structure in adaptive signal. `WZA` combines SNP-level association statistics across genomic windows, whereas the top-candidate method tests whether a region contains an excess of highly associated SNPs. Booker et al. (2024) found that both methods performed as well as or better than single-SNP methods (Kendall’s *τ*, `BayPass`, `LFMM`, and `RDA`) under both strong and weak local adaptation, which is why we use them as our benchmarks here. Comparing SHAP-based attribution from `EcoLocator` to these methods provides a direct test of whether SNP-level importance scores derived from a trained predictive model can recover adaptive loci while preserving resolution at the level of individual variants.

#### Weighted-Z Analysis (WZA)

Following Booker et al. (2024), we first computed per-SNP association statistics by measuring the correlation between genotype and environmental values, and obtained a *p*-value for each SNP. These SNP-level *p*-values were then transformed into Z-scores and combined within genomic windows using a weighted Z-test, where weights are proportional to expected heterozygosity, such that SNPs with higher heterozygosity contribute more strongly to the window-level statistic. This weighting scheme reflects the greater information content of polymorphic sites with intermediate allele frequencies. To account for variation in the number of SNPs per window, we applied a polynomial-based correction in which the expected mean and variance of window-level Z-scores were modeled as functions of SNP count. Corrected *p*-values (Z pVal) were then obtained under the assumption of normality.

#### Top-candidate method

Following Yeaman, Hodgins, et al. (2016), we ranked SNPs genomewide by their association p-values and designated those in the top 1% (i.e. above the 99th percentile) as candidate SNPs. For each window, we then used a one-sided binomial test to assess whther the observed number of candidate SNPs exceeded the number expected by chance, given the window’s total SNP count and the genome-wide candidate rate.

#### Integration across methods

For each simulation replicate, we combined SNP-level SHAP FDR scores, window-level WZA *p*-values, and top-candidate *p*-values into a unified dataset. Window-based statistics were mapped back to individual SNPs using SNP-to-window assignments, allowing direct comparison across methods at the SNP level. To obtain a genome-wide assessment, variant-level results were concatenated across simulation replicates within each genetic architecture and simulation regime.

#### Performance evaluation

Method performance was evaluated using precision–recall (PR) curves, which are appropriate for imbalanced datasets with relatively few adaptive variants. For each method, *p*-values (or, for SHAP, empirical FDR scores) were transformed to scores (*−* log_10_), such that larger values correspond to stronger evidence of association. Because area under the precision–recall curve (AUC-PR) depends only on the rank ordering each score induces, comparing methods this way does not require their underlying significance measures to share a common statistical interpretation or null distribution — only that each score ranks variants consistently from most to least likely adaptive within its own method. Precision and recall were computed across all thresholds, and AUC-PR was used as a summary statistic. The random baseline was defined as the proportion of adaptive variants in the dataset. Performance was evaluated separately within each simulation regime, allowing direct comparison of method behavior under both spatially explicit and Wright–Fisher evolutionary scenarios.

### Empirical application to coastal Douglas-fir

#### Samples and genotyping

Our empirical dataset consisted of open-pollinated seedlings from 348 unrelated coastal Douglas-fir (*Pseudotsuga menziesii* var. *menziesii*) trees sampled across Oregon and Washington. The maternal parents of these trees were woodsrun (unimproved) ‘plus’ trees, identified through field observations as well-adapted to their local environment and exceptional for diameter, height, and stem form. Leaf-derived DNA from individual trees was genotyped using the 58K Douglas-fir Axiom Affymetrix SNP array (Howe et al., 2020). Genotype calls were generated in Axiom Analysis Suite (v.5.4) using a high-stringency workflow (dQC *≥* 0.90, call rate *≥* 0.97), supplemented by a secondary workflow for samples of slightly lower quality (dQC ¿ 0.82, call rate *≥* 0.90). We restricted analysis to SNPs classified as PolyHighResolution — Axiom’s designation for markers with clearly resolved, high-confidence biallelic genotype clusters — yielding 18,732 loci and excluding lower-confidence and off-target variants. Within this marker set, missing genotype rates ranged from approximately 0.15% to 9.85% across samples and were resolved using the imputation procedure described above. The Douglas-fir genome is large and incompletely assembled (*∼*16 Gb; Neale et al., 2017), currently being established by re-sequencing and re-assembly (Amanda de la Torre, pers. comm.). Given this uncertainty, chromosome- or window-level analyses are precluded. Douglas-fir exhibits low self-fertility and seeds derived from maternal parents are typically pollinated by multiple nearby pollen donors. While the pollen pool is heterogeneous, pollen and seed movement in Douglas-fir is limited (typically *<* 400 m), making the maternal source environment a reasonable proxy for the source of open-pollinated seedlings.

#### Climate variables

To represent source climates, we chose 1970–2000 baseline climate normals modeled by ClimateNA (Mahony et al., 2022) using source latitude, longitude, and elevation. We focused on three climate variables shown to be significantly associated with Douglas-fir growth and local adaptation (St. Clair, Mandel, and Vance-Borland, 2005): mean cold month temperature (MCMT), which shows the strongest associations with the timing of summer budset and annual height increment; and Summer Heat:Moisture Index (SHM) and mean summer precipitation (MSP; May–September precipitation), both of which show the strongest associations with the timing of spring bud break and the magnitude of stem taper. These variables capture the primary axes of climate variation across the species’ range and correspond directly to the selection pressures (e.g., winter cold temperatures; aridity) that are managed and mitigated by USFS seed zones. To characterize the spatial scale over which each climate variable varies, we computed a simple spatial correlogram: for each variable, we standardized true values across all 348 sampled trees, calculated the product of standardized values for every pair of trees, and averaged these products within geographic distance bins. We report the approximate distance at which this average first falls below 0.1 as the effective range of spatial autocorrelation for each variable.

#### Model training and feature attribution

Prior to model training, we applied the preprocessing steps described above, and geographic coordinates and environmental covariates were centered and scaled. We trained `EcoLocator` on this dataset using leave-one-out cross-validation, enabling direct comparison between each individual’s predicted and known geographic coordinates and climate values. We also wanted to know whether the climate predictions contain information that the geographic predictions do not. For this we took each predicted location, looked up its climate in ClimateNA, and compared the accuracy of these indirect estimates to the climate values `EcoLocator` predicted directly. SHAP values were summarized across leave-one-out folds by calculating the mean absolute attribution for each of the 18,732 SNPs separately for latitude, longitude, MCMT, MSP, and SHM. For each prediction target, we retained the 179 highest-attribution SNPs, approximately the top 1% of the ranked SHAP distribution, as candidate loci for downstream comparison and biological interpretation. This threshold was used for prioritization rather than as a test of statistical significance. We then compared pairwise overlap among the five target-specific SNP sets to determine the extent to which geographic and climatic predictions relied on shared genomic features. As a supplemental analysis, we aligned array probe sequences to a recent draft assembly (DFv5.29) and, restricting to primary, high-confidence alignments (MAPQ *≥* 30), recovered approximate chromosomal positions for a subset of the genotyped SNPs. We use these positions only to plot attribution scores along the genome; they play no role in prediction.

#### Functional annotation

To characterize the potential biological functions represented among high-attribution loci, SNP identifiers were used to retrieve their source cDNA sequences (Howe et al., 2020). Available sequence annotations included best protein matches to *Arabidopsis thaliana* and *Populus trichocarpa*, along with Mercator gene descriptions and PANTHER functional classes and family identifiers. We used these annotations descriptively to identify recurring biological functions and functional themes among high-attribution loci and to compare these patterns across prediction targets. Because these functional assignments rely in part on homology to angiosperm proteins and SHAP quantifies predictive importance rather than causality, we interpret the resulting annotations as a means of prioritizing and biologically contextualizing candidate loci rather than as evidence that individual SNPs are causal targets of local adaptation.

#### Validation against USFS seed zones

To assess whether prediction errors are biologically meaningful, we compared `EcoLocator`-predicted climate values to the climate ranges within USFS seed transfer zones. In the Pacific west, ‘first-generation’ seed transfer zones (USDA Forest Service, 1973) were narrowly defined to restrict movement and minimize maladaptive responses to winter cold temperatures. ‘Second-generation’ zones (Randall, 1996; Randall and Berrang, 2002) were sub-sequently developed after common garden experiments revealed that Douglas-fir populations show broader climate tolerance than is reflected in first-generation zones. To represent seed zone climate factors (MCMT, MSP, SHM), we used compiled climate factor summaries for first-generation Oregon and Washington seed zones (Shalev et al., 2026). For our analysis, we used 1000-foot elevation band summaries instead of the original 500-foot summaries; this allows us to evaluate results in the spatial context of first-generation zone boundaries (which are widely used in forest management), while allowing comparisons to the broader climate transfers reflected in experimentally-supported second-generation zones. For each individual tree, we compared `EcoLocator` climate predictions to known seed zone climates using two approaches: (a) a “fixed-zone” approach where map-based seed zones were treated as fixed constraints; and (b) a “focal point” approach where the climate range of the local seed zone was used to define the maximum acceptable climate transfer. With the fixed-zone approach, a climate prediction is considered a “match” if it falls within the percentile-defined range of the source seed zone and elevation band. This map-based approach can lead to non-matches when predictions are nearly identical to true values due to the arbitrary nature of fixed map boundaries. With the focal point approach, the difference between the predicted and true climate values is compared to the range of climate variation within the source seed zone. For example, if a source seed zone–elevation band shows a MCMT 100th percentile range of 2°C, an `EcoLocator` MCMT prediction within 2°C of the true value falls inside the range of allowable transfers and is considered a “match”. In both cases we compared predicted values to the 100th percentile range (defined by the minimum and maximum values) and the 98th percentile range (defined by the 1st and 99th percentile values). The indirect location plus ClimateNA estimates were scored the same way.

## Results

### EcoLocator prediction accuracy scales proportionally with strength of local adaptation

We evaluated `EcoLocator` across a range of simulation parameterizations (Figure 2). Following Booker et al. (2024), we ran 100 simulation replicates for each parameter combination in our continuous space simulations, and we ran 30 replicates for the discrete space simulations. We additionally summarized the distribution of local adaptation scores across representative parameter ranges to illustrate how dispersal, selection strength, and genetic architecture influence the degree of local adaptation (see Figure S4). Under restricted dispersal (*σ_D_* = 0.1) and stronger selection (*σ_K_*=0.5), simulated populations became strongly locally adapted to their environments (LA *<* 0.1) after 10,000 generations. `EcoLocator` performed best when dispersal was constrained (*σ_D_*= 0.1, Lon. *R*^2^=0.968, Lat. *R*^2^=0.967, Env. *R*^2^=0.769; Figure 2, panel a), while increasing dispersal reduced accuracy for both geographic and environmental prediction (*σ_D_*=0.5, Lon. *R*^2^=0.815, Lat. *R*^2^=0.814, Env. *R*^2^=0.336; Figure 2, panel a). This pattern is consistent with increasing gene flow weakening both isolation-by-distance structure and genotype–environment correspondence. Weakening selection also reduced environmental prediction accuracy (*σ_K_*=0.3, Env. *R*^2^=0.609; *σ_K_*=0.5, Env. *R*^2^=0.561; *σ_K_*=0.7, Env. *R*^2^=0.421). In contrast, geographic prediction remained largely unchanged across selection regimes (*σ_K_*=0.3, Lon. *R*^2^=0.917, Lat. *R*^2^=0.916; *σ_K_*=0.5, Lon. *R*^2^=0.911, Lat. *R*^2^=0.911; *σ_K_*=0.7, Lon. *R*^2^=0.906, Lat. *R*^2^=0.907). This dissociation is itself informative. Dispersal, and thus the spatial structure of the population, was held constant across selection regimes, and on a landscape with no spatial autocorrelation in the environment (*H* = 0), geography carries no information about the environmental target. The environmental signal that `EcoLocator` recovers here must therefore come from something other than geographic structure.

Varying genetic architecture produced only a slight difference in local adaptation (ΔLA=0.008 between GA1 and GA3), despite the higher number of QTL mutations arising under genetic architecture 3 (GA1=25, GA3=35). Geographic prediction accuracy remained nearly identical across architectures (GA1 - Lon. *R*^2^=0.967, Lat. *R*^2^=0.967; GA2 - Lon. *R*^2^=0.967, Lat. *R*^2^=0.967; GA3 - Lon. *R*^2^=0.968, Lat. *R*^2^=0.968), and environmental covariate prediction accuracy was similarly stable (GA1 - Env. *R*^2^=0.759; GA2 - Env. *R*^2^=0.771; GA3 - Env. *R*^2^=0.787). Although genetic architecture 1 exhibited more extreme outliers and genetic architecture 3 showed slightly higher environmental prediction accuracy, `EcoLocator` performance was relatively robust to the genomic distribution of adaptive loci within the range explored here.

To test whether this robustness extends to landscapes where environmental variation covaries with geography, we ran additional supplemental simulations using genetic architecture 3, under two combined dispersal/selection regimes: limited dispersal with strong selection (*σ_D_* = 0.1, *σ_K_* = 0.5), and higher dispersal with weaker selection (*σ_D_*= 0.5, *σ_K_* = 0.7) (Figure S10). Per-replicate leave-one-out cross validation (LOOCV) *R*^2^ again declined with increasing dispersal and weakening selection under this design, most sharply for latitude (Figure S11). `EcoLocator` also performed strongly under the discrete-space, Wright–Fisher simulation regime (Figure 2, panel b). Across 30 simulation replicates, `EcoLocator` accurately predicted both deme coordinates and the environmental optimum assigned to each deme (Lon. *R*^2^=0.986, Lat. *R*^2^=0.985, Env. *R*^2^=0.916). This is not surprising given how these simulations are built: migration is restricted to neighboring demes, so spatial structure is strong, and local adaptation is driven by a small number of causal loci. The same features account for the stronger attribution and GEA results we report for this regime below. We summarize the full simulation *R*^2^ results in Table S2.

### Feature attribution methods reveal adaptive loci *in silico*

To evaluate whether feature attribution identifies adaptive loci, we summarized SHAP values across all simulation replicates under strong local adaptation (*σ_D_* = 0.1, *σ_K_* = 0.5) and compared attribution scores between non-neutral variants and neutral variants, including sites in linkage disequilibrium. Across all continuous-space genetic architectures, non-neutral variants consistently received higher SHAP values than neutral sites, indicating that `EcoLocator` preferentially attributes predictive signal to loci under selection (Figure 3, panels a–c). This separation was evident both in individual simulation examples and in distributions aggregated across all replicates.

**Figure 3:**
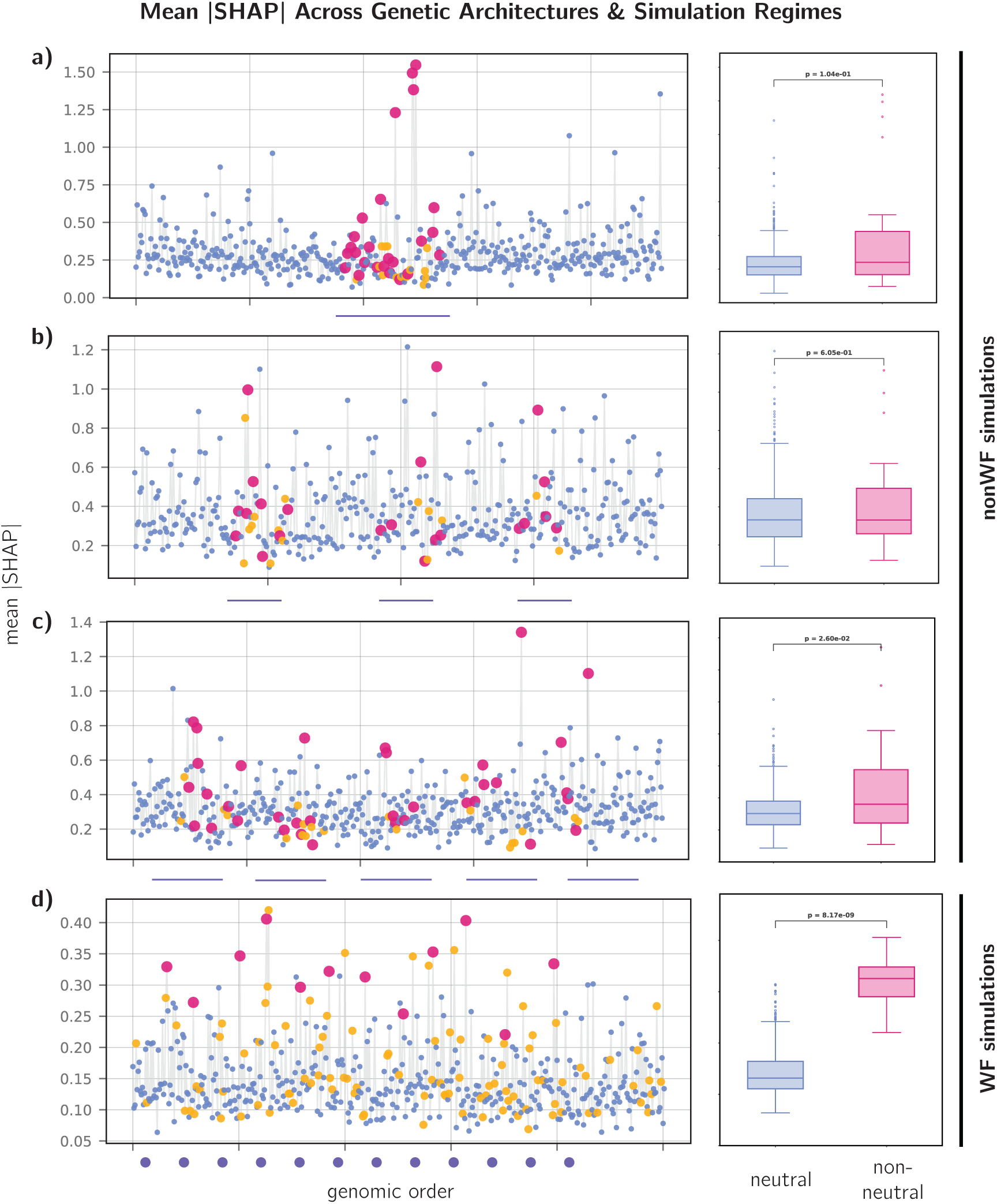
Feature attribution across genetic architectures. Each panel on the left shows mean absolute SHAP scores across the simulated chromosome for neutral sites (blue), non-neutral sites (pink), and sites in linkage disequilibrium with non-neutral variants (yellow). **Panel a)** represents simulations under genetic architecture 1, **Panel b)** represents simulations under genetic architecture 2, and **Panel c)** represents simulations under genetic architecture 3. Across panels a, b, and c, dispersal was held constant at (*σ_D_* = 0.1), and the standard deviation of the fitness function was held constant at (*σ_K_* = 0.5). In panels a, b, c, horizontal lines in the x axis indicate the location of the causal loci. **Panel d)** shows SHAP attributions for the Wright–Fisher, discrete-space simulation regime, where 12 large-effect causal loci (shown as dots on the x-axis) were simulated. For all simulations, bloxplots on the right show the distribution of absolute SHAP values between neutral and non-neutral sites.

Under genetic architecture 1 (Figure 3, panel a), non-neutral variants exhibited significantly higher mean attribution values (*µ* = 0.390) relative to neutral and LD sites combined (*µ* = 0.292; *p* = 2.21 *×* 10*^−^*^4^). A similar pattern was observed under genetic architecture 2 (Figure 3, panel b), where non-neutral variants again showed elevated SHAP values (*µ* = 0.403) compared to neutral and LD variants (*µ* = 0.303; *p* = 5.85 *×* 10*^−^*^6^). Despite differences in the number of adaptive loci, genetic architecture 3 (Figure 3, panel c) also showed enrichment of SHAP signal at non-neutral variants (*µ* = 0.394) relative to neutral and LD sites (*µ* = 0.319; *p* = 8.79 *×* 10*^−^*^4^).

Although effect sizes were modest (Cohen’s *d ≈* 0.14–0.18 across architectures), the consistent shift in the distribution of attribution values indicates that adaptive loci are preferentially weighted by the model. Notably, the magnitude of separation between non-neutral and neutral variants was similar across genetic architectures, despite differences in the total number of causal loci and the extent of linkage disequilibrium. This observation suggests that feature attribution performance is robust to variation in genetic architecture within the range explored here, which also held under the spatially correlated environment design described above (Figure S10): non-neutral loci again showed elevated attribution across all three targets, and longitude and environment tracked the same genomic regions in a representative replicate (Figure S12), an expected consequence of the two variables following a shared spatial gradient by design. We observed stronger separation between neutral and non-neutral sites in the discrete-space Wright–Fisher simulations (Figure 3, panel d), where local adaptation was driven by 12 causal loci. Non-neutral variants had substantially higher SHAP values than neutral variants, with a median attribution value of 0.326 for non-neutral sites compared to 0.132 for neutral sites. This difference was highly significant (*p* = 8.17 *×* 10*^−^*^9^), indicating that `EcoLocator` concentrated attribution signal at causal loci in this discrete-deme simulation regime. These results show that SHAP-based feature attribution can highlight loci contributing to local adaptation, while also capturing signal from linked regions.

Permutation-based sensitivity analyses further showed that predictive signal was not restricted to the selected QTL window. Permuting the selected window alone produced only modest changes in prediction performance, whereas jointly permuting the selected window and all LD-linked sites caused accuracy to fall below zero for all three targets (X *R*^2^ = *−*0.114, Y *R*^2^ = *−*0.181, Env. *R*^2^ = *−*0.267; Figure S3). Because these *R*^2^ values are computed out-of-sample on permuted genotypes, they can be negative: a negative value means the model’s predictions are worse than simply guessing the training-set mean for every individual, indicating that little or no usable signal remains once those sites are disrupted. Random genome-wide permutation produced a more gradual decline in performance, with complete randomization similarly eliminating predictive accuracy (X *R*^2^ = *−*0.507, Y *R*^2^ = *−*0.426, Env. *R*^2^ = *−*0.425).

### Comparison of EcoLocator to other GEA methods

A notable capability of machine learning approaches like `EcoLocator` is that, in this context, feature attribution methods provide locus-level importance scores that can be interpreted analogously to genome–environment association statistics. This allows direct comparison between model-derived attribution scores and traditional GEA approaches for identifying adaptive loci. Using SHAP values computed from trained `EcoLocator` models, we ranked variants according to their contribution to environmental prediction and compared their performance to Weighted-Z Analysis (`WZA`) (Booker et al., 2024) and the top-candidate method (Yeaman, Hodgins, et al., 2016).

To quantify detection accuracy, we calculated empirical FDR scores from SHAP rankings, calibrated against known true adaptive status rather than a null distribution (see Methods), and evaluated performance using precision–recall curves (Figure 4). We refer to this `EcoLocator`-plus-SHAP pipeline as `EcoLocator` ×SHAP below, since SHAP attribution is what turns `EcoLocator`’s trained model into a method that can be benchmarked directly against GEA approaches like `WZA` and top-candidate. The random baseline represents the AUC-PR expected if variants were ranked at random, given the proportion of adaptive variants in each simulation regime. For each simulation regime, the precision–recall curves were constructed by pooling SNPs across all simulation replicates, so the resulting AUC-PR values reflect global discriminatory performance. We also calculated replicate-level AUC-PR values, shown as violin plots. Thus, the pooled AUC-PR values summarize overall variant-level discrimination across all replicates, whereas the violin plots summarize replicate-to-replicate variation in method performance.

**Figure 4:**
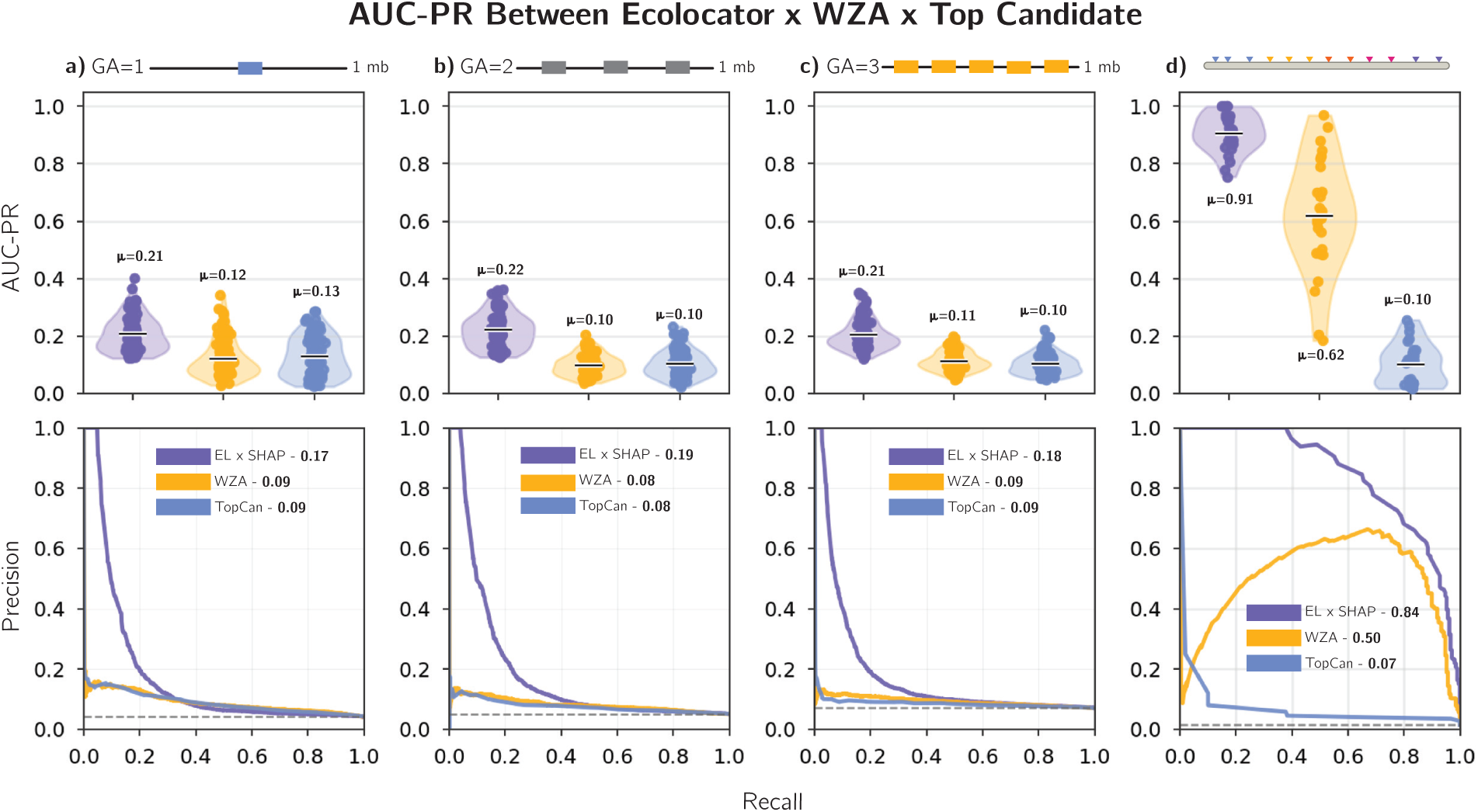
AUC-PR curve comparison between SHAP, `WZA`, and top-candidate methods. Each column represents a different simulation regime. **Column a)** represents genetic architecture 1, **Column b)** represents genetic architecture 2, and **Column c)** represents genetic architecture 3. **Column d)** represents the Wright–Fisher, discrete-space simulation regime. The top row shows violin plots of per-replicate AUC-PR values, where each point represents one simulation replicate, the horizontal bar indicates the mean, and the annotated value gives the mean AUC-PR for each method; this view captures replicate-to-replicate variability in performance. The bottom row shows precision-recall curves constructed by pooling SNPs across all simulation replicates within each regime, with the annotated AUC-PR giving the area under each pooled curve; this reflects global discriminatory performance and is not directly comparable to the per-replicate means shown above. In all panels, purple indicates `EcoLocator` ×SHAP, yellow indicates `WZA`, and blue indicates the top-candidate method.

Across all continuous-space genetic architectures, SHAP-based attribution consistently recovered a higher proportion of true adaptive variants than either comparison method in the pooled precision–recall analysis. Under genetic architecture 1, SHAP achieved a pooled AUC-PR of 0.1734, substantially exceeding `WZA` (0.0911) and the top-candidate approach (0.0890), and outperforming the random expectation (0.0426). Under genetic architecture 2, SHAP again showed improved performance (pooled AUC-PR = 0.1870) relative to `WZA` (0.0832) and the top-candidate method (0.0767), with a random baseline of 0.0514. Similarly, under genetic architecture 3, SHAP yielded a pooled AUC-PR of 0.1786, compared to 0.0946 for `WZA` and 0.0874 for the top-candidate approach (random = 0.0713). The replicate-level violin plots showed the same overall ranking. Under genetic architecture 1, `EcoLocator` ×SHAP had the highest mean AUC-PR (mean AUC-PR = 0.2076), followed by the top-candidate method (mean AUC-PR = 0.1281) and `WZA` (mean AUC-PR = 0.1212). Under genetic architecture 2, `EcoLocator` ×SHAP again showed the highest mean performance (mean AUC-PR = 0.2211), followed by the top-candidate method (mean AUC-PR = 0.1032) and `WZA` (mean AUC-PR = 0.0995). Under genetic architecture 3, `EcoLocator` ×SHAP maintained the highest mean AUC-PR (mean AUC-PR = 0.2054), followed by `WZA` (mean AUC-PR = 0.1133) and the top-candidate method (mean AUC-PR = 0.1049). Performance differences were most pronounced at low recall values, where SHAP maintained substantially higher precision, indicating that the highest-ranked loci were enriched for true causal variants. In contrast, `WZA` and the top-candidate method showed rapid declines in precision, approaching random expectations across much of the recall range. Notably, SHAP maintained improved performance across all genetic architectures, despite differences in the number and genomic distribution of adaptive loci. SHAP-based attribution retained its advantage over `WZA` and the top-candidate method under this same spatially correlated design (Figure S8). In the discrete-space Wright–Fisher simulations, `EcoLocator` ×SHAP showed particularly strong recovery of adaptive loci, achieving a pooled AUC-PR of 0.8430 relative to a random baseline of 0.0149 (Figure 4, column d). `WZA` also performed well in this regime (pooled AUC-PR = 0.4983), whereas the top-candidate method showed substantially lower recovery (pooled AUC-PR = 0.0714). Replicate-level AUC-PR values followed the same ordering, with `EcoLocator` ×SHAP showing the highest mean performance (mean AUC-PR = 0.9061), followed by `WZA` (mean AUC-PR = 0.6191) and the top-candidate method (mean AUC-PR = 0.1017).

### Predicting suitable environments for coastal Douglas-fir using EcoLocator

Simulations provide a controlled setting to benchmark `EcoLocator`, but the ultimate goal is to evaluate its performance in natural systems where demography, gene flow, selection, and management history interact in complex ways. To this end, we applied `EcoLocator` to 348 coastal Douglas-fir trees sampled across Oregon and Washington (Figure 5, panel f) and genotyped at 18,732 SNPs. For each tree we predicted geographic origin along with three climate variables (MCMT, SHM, and MSP) known to structure local adaptation in this species, using leave-one-out cross-validation so that every prediction could be compared to a known value.

**Figure 5:**
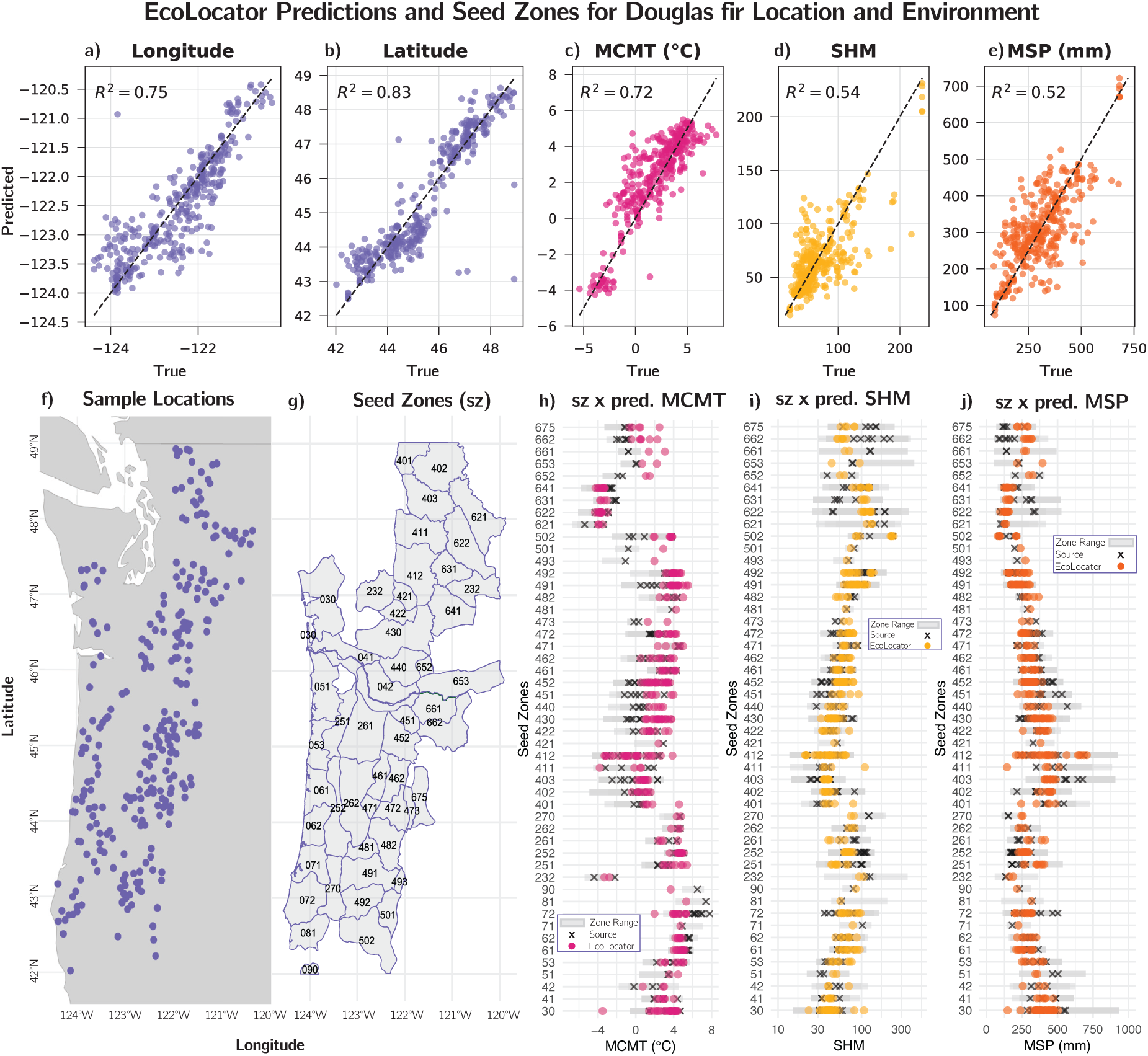
Douglas-fir case study results. **Panels a-e.** show true vs. predicted scatterplots across all prediction targets. From left to right, we show Longitude (*R*^2^=0.75), Latitude (*R*^2^= 0.83), Mean Cold Month Temperature (MCMT; *R*^2^= 0.72), Summer Heat:Moisture Index (SHM; *R*^2^= 0.54), and Mean Summer Precipitation (MSP; *R*^2^= 0.52). **Panels f-g.** both show the sampling range, with **Panel f.** being the sample locations and **Panel g.** being the seed zone map, with labels only on the seed zones containing samples. **Panels h-j.** show where `EcoLocator` predictions fall within the established seed zones, with gray boxes showing the environmental covariate’s ranges in a given zone, ’*×*’s’ representing the source sample value, and colored dots representing `EcoLocator` predictions inside each zone. From left to right, we show MCMT in pink, SHM in yellow, and MSP in orange.

#### Prediction accuracy

`EcoLocator` predicted source latitudes and longitudes with *R*^2^ values of 0.83 and 0.75, respectively (Figure 5). Combined latitude and longitude for each sample gave predicted geographic origins that showed a median error of 51.6km (see Figure S6), with minimum errors as low as 1.2km and maximum errors as high as 646km. Errors greater than 200km appeared as extreme outliers and were not associated with obvious geographic (e.g., edge of range; clustered locations) or genetic (e.g., unusual homozygosity) attributes. Label errors or failed predictions may contribute to these unusually large errors.

Climate is strongly spatially autocorrelated, so one might worry that the climate predictions simply ride on the geographic ones. If that were the case, predicting location and then looking up its climate in ClimateNA would do at least as well as predicting climate directly. It does not. Direct estimates from `EcoLocator` were more accurate than ClimateNA values at predicted locations for all three variables (paired Wilcoxon signed-rank test on absolute error, *n* = 348; MCMT *p* = 1.1*×* 10*^−^*^6^, MSP *p* = 6.0 *×* 10*^−^*^10^, SHM *p* = 1.6 *×* 10*^−^*^13^; Table S1), so the model is picking up climate-related signal in the genotypes that geography alone does not supply. Among the three modeled climate variables, Mean Cold Month Temperature (MCMT) showed the highest predicted *R*^2^ of 0.72, with Mean Summer Precipitation (MSP) (*R*^2^=0.52) and Summer Heat:Moisture Index (SHM) (*R*^2^=0.54) showing lower and comparable prediction accuracies. This pattern is consistent with differences in the spatial scale of each variable’s underlying gradient: across our sampled trees, MCMT remains spatially autocorrelated out to roughly 150–175 km, whereas MSP and SHM decorrelate by around 100–125 km, likely reflecting the influence of local topography (e.g., rain shadows and orographic effects) on precipitation-related variables (Frankson et al., 2022). Because `EcoLocator`’s genomic signal is itself spatially structured, climate variables that vary smoothly across the landscape should be easier to predict from that structure than variables that vary sharply over short distances.

#### Validation against USFS seed zones

To assess whether `EcoLocator`’s prediction errors are biologically meaningful, we compared `EcoLocator`-predicted values to the climate ranges within USFS seed zone–elevation bands, using the fixed-zone and focal point criteria described in Methods. Using the fixed-zone approach, we find that 66.1% of the MCMT predictions fall within seed zone–elevation band 100th percentile ranges, and that the number of matches drops to 49.7% of predictions when using the 98th percentile range. The proportion of matches is higher for MSP (86.2% for the 100th percentile range) and SHM (87.6% for the 100th percentile range) (Table 1). These values are all larger than those estimated using ClimateNA predictions from `EcoLocator` geographic predictions (Table 1).

**Table 1:** Comparison of EcoLocator and Location+ClimateNA predictions across climate variables using the fixed-boundary seed zones.

| Factor | EcoLocator |  | Location+ClimateNA |  |
| --- | --- | --- | --- | --- |
|  | 100%ile range | 98%ile range | 100%ile range | 98%ile range |
| MCMT | 66.1% | 49.7% | 65.2% | 48.9% |
| MSP | 86.2% | 75.9% | 74.1% | 57.5% |
| SHM | 87.6% | 74.7% | 71.6% | 55.5% |

Using the focal point approach, we find that 93.1% of the MCMT predictions show lower deviations from the true value than the 100th percentile range of the source seed zones. The number of matches drops to 87.1% of predictions when using the 98th percentile range (Table 2). Predictions for MSP and SHM showed higher accuracies, with 98.6% and 97.7% of estimates falling within the 100th percentile range of the source seed zone, respectively. As was observed with predictions within seed zones, `EcoLocator` showed higher prediction accuracy than estimates using ClimateNA predictions from `EcoLocator` geographic predictions (Table 2).

**Table 2:** Comparison of EcoLocator and Location+ClimateNA predictions across climate variables using the focal point seed zones.

| Factor | EcoLocator |  | Location+ClimateNA |  |
| --- | --- | --- | --- | --- |
|  | 100%ile range | 98%ile range | 100%ile range | 98%ile range |
| MCMT | 93.1% | 87.1% | 89.9% | 75.9% |
| MSP | 98.6% | 93.1% | 94.5% | 85.3% |
| SHM | 97.7% | 92.8% | 91.9% | 83.6% |

Prediction error for different climate variables was not uniform across seed zones, and patterns differed by MCMT, MSP, and SHM (Figure 5, panels h-j). In our sample, trees from coastal seed zones in Oregon and Washington (e.g., zones 031, 041, 051, 061) showed lower prediction errors for MCMT than either Cascade or Cascade Crest sources. Conversely, trees from Oregon and Washington Cascades seed zones showed lower prediction errors for MSP and SHM than did Coastal or Cascade Crest trees. These results demonstrate that `EcoLocator` can predict the climate of origin of individual genotypes at a resolution finer than the climatic breadth of existing seed zones, the tolerance that actually governs seed transfer decisions. Our results further illustrate the value of seed zone benchmarking as a biologically grounded validation framework: predictions that fall within a seed zone climate envelope are not just numerically accurate, but are close enough that forest managers consider it an acceptable transfer source.

Although the genotype array used for inference carries no genomic coordinates, our alignment to the DFv5.29 draft assembly placed a subset of SNPs on chromosomes, and plotting mean *|*SHAP*|* against these positions lets us view attribution scores in genomic context for each of the five prediction targets (Figure S9). Visually, several genomic regions of Figure S9 show elevated attribution for both longitude and MCMT simultaneously, consistent with the fact that Mean Cold Month Temperature covaries with longitude across coastal Douglas-fir’s range, with coastal populations experiencing milder winters than those toward the Cascade crest—the same pattern we recover in simulation when environment is spatially correlated with geography (Figure S10, Figure S12).

We next asked whether the same genomic features contributed strongly across geographic and climatic prediction targets. Comparing the 179 highest-attribution SNPs for each target revealed substantial variation in pairwise overlap (Table S3). Only 13 SNPs were shared between latitude and MCMT (7.3%), whereas longitude and MCMT shared 97 SNPs (54.2%). The strongest overlap occurred between MSP and SHM, which shared 133 SNPs (74.3%). These overlap patterns mirror relationships among the prediction targets themselves. MSP and SHM both capture variation related to summer moisture availability, a relationship that is reflected by the largest overlap among high-attribution SNPs. Similarly, MCMT varies strongly along the longitudinal transition from relatively mild maritime environments toward colder environments closer to the Cascade Range and shares substantial genomic signal with longitude. In contrast, latitude and MCMT shared relatively few high-attribution loci (7.3%), consistent with latitude representing a north–south geographic axis rather than the longitudinal winter-temperature gradient captured by MCMT. Because MSP and SHM are themselves strongly correlated (SHM is calculated in part from MSP) and MCMT covaries with longitude across the sampled range, some of this overlap likely reflects intercorrelation among the targets themselves rather than independent biological signal specific to each; the overlap patterns are informative about which loci the model draws on for correlated targets, not evidence that those loci are independently causal for both.

Functional annotation of the high-attribution SNPs revealed additional differences among prediction targets. Because Axiom array probes are designed from cDNA sequence (see Methods), each genotyped SNP already has a known transcript-level origin, so “loci” here refers to SNPs anchored within an expressed gene’s transcript sequence, rather than an anonymous genomic position matched to the nearest annotated gene. Among loci with informative annotations, latitude-associated candidates included putative homologs involved in phytochrome-mediated light signaling, circadian and seasonal regulation, and developmental timing, including *PIF3*, *FPA*, WNK-family kinases, and several MYB transcription factors. MCMT- and longitude-associated candidates included putative *ELF3* and DREB/AP2-family transcription factors, SPL-family regulators, molecular chaperones including DnaJ proteins, membrane lipid and desaturase-associated genes, and genes involved in carbohydrate and trehalose metabolism. These annotations represented functions associated with seasonal regulation, temperature-stress response, membrane remodeling, and acclimation (C.-M. Lee and Thomashow, 2012; Chen and Thelen, 2013).

A distinct but internally consistent set of functions was observed among the strongly overlapping MSP and SHM candidate sets. These included putative PYL-family ABA receptors, PP2C phosphatases (Park et al., 2009), LEA proteins, an osmosis-responsive factor, GST-family proteins, membrane transporters, and other genes associated with cellular stress protection. Additional candidates across the MSP and SHM sets were associated with heat-shock, mitochondrial stress, and redox-response functions. Although many high-attribution loci were poorly annotated or had functions that were difficult to relate directly to climate, the interpretable candidates repeatedly represented processes associated with environmental timing, temperature acclimation, water-stress signaling, membrane remodeling, and oxidative-stress response. The complete set of high-attribution SNPs and associated functional annotations is provided in Supplementary Data S1.

## Discussion

### Joint prediction of geography and climate reveals spatial structure and local adaptation

It is well established that multilocus genomic variation encodes both neutral spatial structure and signals of environmental adaptation. Genetic principal components mirror geography (Novembre, Johnson, et al., 2008), allele frequencies covary with environmental gradients under local selection (Coop et al., 2010; Savolainen, Lascoux, and Merilä, 2013), and isolation by environment can generate genetic structure independent of geographic distance (Wang and Bradburd, 2014). Continuous spatial structure leaves particularly strong and pervasive signatures in genomic data, influencing diversity statistics, demographic inference, and genotype–environment associations in ways that discrete-deme models fail to capture (Battey, Ralph, and Kern, 2020b; Chevy et al., 2025). The method `Locator` (Battey, Ralph, and Kern, 2020a) demonstrated that these spatial signatures are rich enough to predict the geographic origin of individuals from genotypes alone using deep neural networks, achieving high accuracy across diverse taxa. `EcoLocator` extends this framework by jointly predicting geographic location and climate of origin, unifying spatial and environmental inference in a single supervised model. This allows us to ask not only where an individual comes from, but what environmental conditions it is adapted to, and how the recovery of each signal depends on evolutionary parameters. In our simulations, increasing dispersal hurt both geographic and environmental prediction, but weakening selection hurt only environmental prediction. This decoupling is consistent with the expectation that these two signals have distinct evolutionary origins: geographic prediction depends on isolation-by-distance patterns driven by limited gene flow, whereas environmental prediction requires allele frequency clines that track ecological gradients—a signal that is eroded when gene flow overwhelms local selection. Prediction accuracy barely changed across the three genetic architectures we tested, though we note that these architectures cover only a small part of the genetic complexity expected in natural populations.

Our primary continuous-space simulations deliberately decoupled environment from geography by imposing zero spatial autocorrelation on the environmental layer (*H* = 0), a much sharper decoupling than real landscapes typically show: in our Douglas-fir dataset, climate variables remain spatially autocorrelated out to roughly 100–175 km (see above), whereas environmental gradients commonly covary with geographic structure in natural landscapes more generally (Meirmans, 2012). We therefore repeated our analysis under a spatially correlated environmental gradient to ask whether the same general patterns persist when geography and environment share spatial structure. Prediction accuracy again declined under higher dispersal and weaker selection, indicating that the relationships observed in the decoupled simulations persist under this more realistic spatial configuration. However, when geography and environment covary, shared predictive signal cannot necessarily be attributed uniquely to demographic structure or local adaptation, an important limitation that we return to below.

The Wright–Fisher results show what `EcoLocator` can do when the problem is easy. Discrete demes, migration restricted to nearest neighbors, and directional selection on a dozen causal loci give the model a clearly structured task, and relative to the size of the simulated grid, migration is slow enough that spatial and genotype–environment structure persist. In our continuous-space simulations, by contrast, diffuse genetic architectures, high dispersal, or weak selection erode these signals; because dispersal in these simulations is a non-trivial fraction of the landscape width per generation (2–10%, see Methods), individuals mix across the range more readily than in the Wright–Fisher grid. It is likely this relative scale of dispersal to range size, rather than continuous space *per se*, that drives the difference, though we have not tested whether a proportionally larger continuous-space landscape would recover Wright–Fisher-like performance. The contrast between the two regimes is nonetheless a reminder that the difficulty of the prediction problem depends on the underlying biology (Booker et al., 2024; Dauphin et al., 2023).

The Douglas-fir case study extends these findings to a natural system where the ground truth is not fully known. `EcoLocator`’s predictions of geographic origin and climate variables in coastal Douglas-fir are consistent with the species’ well-documented patterns of local adaptation along steep latitudinal and elevational gradients in western North America (Aitken and Whitlock, 2013). That the model performs well in this system despite low population differentiation (Krutovsky et al., 2009), incomplete genomic resources (Neale et al., 2017), and a relatively modest number of SNPs, suggests that the joint prediction framework can recover informative genomic signal even under challenging empirical conditions. Feature attribution provides additional resolution into how this predictive signal is distributed across geographic and environmental targets, which we discuss in detail below.

### Feature attribution as a tool for identifying candidate adaptive loci

SHAP-based feature attribution provides a window into which genomic regions drive `EcoLocator`’s predictions. That `EcoLocator`’s direct climate predictions outperformed a geography-then-ClimateNA lookup in Douglas-fir (see above) already indicates that this signal reflects genuine genotype–environment covariance rather than geography alone; feature attribution lets us ask which loci carry that signal. Our simulation results demonstrate that non-neutral variants consistently receive higher attribution scores than neutral sites, confirming that the model preferentially weights loci under selection. This signal persists across genetic architectures despite modest effect sizes, indicating that SHAP can detect the distributed signature of polygenic adaptation even when individual locus effects are small.

However, the modest effect sizes also highlight an inherent challenge: under polygenic adaptation, the signal at any single locus is subtle, and attribution scores for adaptive and neutral loci overlap substantially. This is consistent with the broader finding that GEA methods are generally underpowered under weak selection, and that detecting most causal loci requires tolerating a substantial false positive rate (Lotterhos and Whitlock, 2015; Booker et al., 2024). `EcoLocator`’s feature attribution faces the same trade-off: thresholding on SHAP values to identify candidate loci will inevitably involve a balance between sensitivity and specificity that depends on the unknown strength of selection and the genetic architecture of adaptation.

The continuous-space simulations therefore represent a comparatively diffuse attribution problem, with adaptive effects distributed across loci and additional predictive signal extending into linked sites through hitchhiking. Linkage disequilibrium further complicates interpretation, as SHAP values are elevated not only at causal variants but also at linked neutral sites. This is expected, because LD spreads adaptive signal across genomic neighborhoods. The recombination-rate comparison illustrates this effect directly: LD extended much more broadly beyond the selected QTL window under 10*^−^*^8^ than under the 10*^−^*^6^ rate used for the analyses reported here (Figure S2). Together, distributed adaptive effects and signal shared among linked sites help explain both the modest separation between neutral and non-neutral SHAP distributions and the difficulty of resolving individual causal sites.

The permutation-based sensitivity analyses complement SHAP by providing a different lens on the same question. By disrupting specific genomic regions and measuring the resulting loss in prediction accuracy, these analyses show how much predictive information is carried by the selected window, by linked sites, and by genome-wide background variation. Prediction declined much more strongly when the selected window and LD-linked sites were permuted together than when the selected window alone was disrupted (Figure S3). These findings suggest that `EcoLocator` is not relying on the selected window alone, but on a broader LD-associated haplotypic signal that can obscure the distinction between causal and linked variants.

Taken together, these results suggest that SHAP is most reliable for identifying candidate genomic regions or haplotypic blocks associated with local adaptation, rather than for pinpointing individual causal mutations. This distinction matters even more in empirical systems, where LD can reflect physical linkage, demographic history, selection, population structure, or unobserved ancestry, and where the true causal variants are rarely known.

Notably, SHAP-based attribution from `EcoLocator` shows greater power to identify adaptive loci than `WZA` (Booker et al., 2024) and top-candidate approaches in our simulation comparisons (Figure 4). This advantage likely reflects the fact that SHAP attribution operates on a model that has already learned the joint relationship between genotypes, geography, and environment, rather than testing each locus independently against a single environmental predictor. By conditioning on a trained multitask model, SHAP can capture contributions from loci whose individual marginal correlations with environment are weak but whose combined effect is detectable in the context of the full genotype. This shows that supervised prediction followed by *post hoc* feature attribution can be a powerful complement to traditional GEA approaches, particularly under polygenic adaptation where per-locus signals are subtle. At the same time, the stronger performance of both SHAP and `WZA` in the Wright–Fisher simulations suggests that locus-identification performance can improve when adaptive signal is strong and localized, whereas more diffuse adaptive architectures present a more difficult inference problem.

One caveat becomes important when feature attribution moves from simulations to real landscapes. Our continuous-space simulations use an environmental layer with no spatial autocorrelation (*H* = 0), so that environment and geographic structure share as little signal as possible. In nature the two usually covary, and when they do, attribution alone cannot tell us whether a high-attribution locus reflects environmental selection, demographic structure, linkage to a selected locus, or some mix of these (we return to this under Strengths and limitations)). High-attribution loci should therefore be read as candidate genomic regions associated with the predicted gradient, not as confirmed targets of adaptation.

The Douglas-fir results nevertheless tell us something about how this signal is structured. The high-attribution SNP sets overlapped most for pairs of targets that track related gradients, MSP with SHM and longitude with MCMT, and hardly at all for latitude and MCMT (Table S3). As noted above, some of this overlap likely reflects correlation among the target variables themselves rather than independently causal shared loci. Still, the low overlap between the more weakly correlated targets (latitude and MCMT, 7.3%) suggests that `EcoLocator` is not simply handing the same spatially structured loci high attribution for every target, and that a meaningful part of the signal is specific to each axis of geographic or environmental variation. The functional annotations echo this pattern. Rather than all five targets converging on the same set of candidate genes, each target’s high-attribution candidates are dominated by a distinct functional theme, and these themes line up with what is already known in coastal Douglas-fir. Latitude candidates were dominated by genes involved in light sensing and seasonal developmental regulation, which fits the known effects of photoperiod and temperature on growth cessation (Ford, Harrington, and St. Clair, 2017). Longitude and MCMT candidates were dominated by genes involved in seasonal regulation and temperature acclimation, in line with genomic evidence for polygenic cold adaptation in this species (De La Torre et al., 2021). MSP and SHM candidates repeatedly involved ABA-mediated water-stress signaling, osmotic protection, and redox response, which matches common-garden evidence for genetic variation in drought and cold hardiness along summer-precipitation and winter-temperature gradients (Bansal, Harrington, and St. Clair, 2016). Viewed together, these patterns suggest that environmental timing, including the timing of growth, dormancy, acclimation, and stress response, may be an important component of the genotype–environment signal captured by `EcoLocator` in Douglas-fir.

### Implications for Douglas-fir management and forestry

For natural resource management, `EcoLocator` addresses a practical need: identifying the geographic and climatic source of biological materials under historical, current, or projected future climates. Seed transfer guidelines have been used on U.S. Forest Service lands for over a half century (Kummel, Rindt, and Munger, 1944), and comparable guidelines extend back nearly a century in Europe (Myking et al., 2016). Despite the history and ubiquity of seed transfer zones, large-scale restoration efforts with non-local materials have been a common occurrence in forestry, both in the management of native forests and in introduced species trials. These unintended ‘local adaptation experiments’ provide clear examples of maladaptive responses, but they are usually associated with a poor understanding of the seed source. Tools like `EcoLocator` can provide estimates of the source location and environment, making it possible to interpret maladaptive responses in large-scale plantings in light of geographic or climatic transfer distances. Applying `EcoLocator` to trials of suspected ‘off-site’ origin could substantially expand our knowledge of growth and fitness responses of large climate transfers, as designed provenance tests with ‘off-site’ materials are limited in number for most tree species.

Similarly, forest tree provenance tests established outside native ranges typically have the goal of identifying high-performing genotypes and geographic sources where additional material can be selected. Incomplete records or missing tree identities limit the utility of these tests by making it difficult to match similar climates to identify pre-adapted seed for screening and breeding. Efforts to combat illegal logging now increasingly rely on DNA-based methods to identify the geographic source of tissues, such as wood. Adding climatic source information to these investigations can help investigators discriminate among geographically similar but climatically distinct sources (such as those separated by elevation). Finally, traditional reforestation programs based on seed transfer guidelines and common-garden trials are increasingly strained by the pace of climate change (Aitken and Bemmels, 2016), and new tools are needed to manage climate uncertainty. Genomic tools that predict the climate of origin of a given genotype offer a unique strategy for answering these questions, enabling managers to evaluate whether replanting resources are well-matched to the conditions they will encounter at a given site.

Coastal Douglas-fir is a natural test case for this application. Its economic importance, extensive provenance trial data, and strong signals of local adaptation along climate gradients provide both the motivation and the validation framework for genomic prediction tools. By predicting climate of origin alongside geographic location, `EcoLocator` provides information that is directly relevant to seed transfer decisions: two seed sources from similar geographic locations may differ in their predicted climate of origin, reflecting adaptive differences that might be missed by geography-based guidelines alone. By comparing `EcoLocator` predictions to Douglas-fir seed zones, these predictions become interpretable in definable management terms. For example, the most recently proposed seed zone–elevation bands in Oregon and Washington span approximately 2.5°C in MCMT across the 98th percentile range (Shalev et al., 2026). Our results suggest that `EcoLocator` predictions fall within these climate tolerances 87.1% of the time with Douglas-fir. Importantly, this level of accuracy was achieved with a modestly sized dataset of 348 trees and approximately 18,000 SNPs, and without any use of genomic position. `EcoLocator` operates on the genotype matrix alone, so neither a contiguous assembly nor an ordered marker map is required—an advantage in species whose genomic resources remain incomplete. Although additional sampling is not evaluated empirically here, downsampling in our simulations (see Figure S5) and previous work with Locator suggest that geographic prediction accuracy increases with increased sampling density and, to a lesser extent, increased genomic sampling (Battey, Ralph, and Kern, 2020a). Similar improvements should be possible for `EcoLocator`, particularly in regions where current predictions are limited by sparse sampling or rapidly changing environmental gradients.

At the same time, the interpretation of `EcoLocator` predictions depends on the sampling history and biological structure of the material being analysed. The Douglas-fir individuals used here were designated as “plus” trees, meaning they were identified through provenance trials or field observations as being particularly well-adapted to their local environments. This selection criterion may strengthen genotype–environment correspondence and aid climate prediction, but it could also overestimate performance relative to less-characterized or more heterogeneous samples. The spatial distribution of prediction error highlights important considerations for applying `EcoLocator` in complex landscapes. For example, the highest prediction errors for MCMT in our dataset were concentrated near the eastern crest of the Cascade Range. Notably, within this crest-adjacent zone, error was significantly higher in the southern Cascades than in the northern Cascades (mean *|*error*|* 1.57*^◦^*C vs. 1.03*^◦^*C; Mann–Whitney *p* = 2.7 *×* 10*^−^*^4^), the opposite of what would be expected if steep, complex terrain alone were driving these errors, since the northern Cascades are topographically steeper (Haugerud and Tabor, 2009). This pattern is also counter to an introgression-based explanation. Evidence of admixture with interior Douglas-fir (*Pseudotsuga menziesii* var. *glauca*) has been described in Washington east of the Cascade crest (St. Clair, Mandel, and Vance-Borland, 2005; Peĺaez et al., 2024), whereas trees in the southern Oregon Cascades—where error is higher— show no evidence of introgression. We do not have a confident explanation for this asymmetry. A possibility is that it reflects post-glacial colonization history, as northern *glauca*-introgressed zones were deglaciated more recently than the southern Cascades, and reduced ancestral genetic variation associated with a more recent colonization history could affect prediction accuracy. Steep elevational gradients and low local sampling density near the crest, and positional inaccuracies in older Township–Range–Section seed sources, may also contribute to reduced prediction accuracy in this region. These patterns suggest that prediction errors point to biologically meaningful features of the study system, including population history, complex topography, and uncertainty in historical source records (Rehmann, Ralph, and Kern, 2024).

### Strengths and limitations

A key strength of `EcoLocator` is its multitask design, which simultaneously models geographic and environmental variation. This joint framework allows the model to learn both shared and target-specific components of genomic variation and, when sufficient information is present in the genotype data, to distinguish signals associated with geographic structure from those associated with environmental variation. This design also bears on a well-documented challenge of conventional GEA methods. When population structure aligns with environmental gradients, as it often does in nature, methods that correct for structure can inadvertently remove true adaptive signal along with neutral confounders (Lotterhos, 2019; Booker et al., 2024). `EcoLocator` takes a different route: rather than regressing geographic structure out, it models geographic and environmental variation as related but distinct prediction tasks. However, this does not resolve the confounding. When climate covaries with geography, non-causal but spatially structured loci can still receive high attribution for climate simply because they predict location, and `EcoLocator` does not attempt to ask whether a given locus carries environmental signal beyond what spatial structure explains, as more classic GEA methods do. Attribution from `EcoLocator` is best interpreted as identifying loci that are informative for the joint prediction task, with the question of which of those are adaptive left to the follow-up analyses discussed above.

Furthermore, the multitask framework is naturally suited to systems where adaptation occurs along multiple environmental axes. Booker et al. (2024) noted that window-based GEA methods can lose power when adaptation involves semi-independent environmental gradients, because the LD patterns around causal alleles may differ across gradients. `EcoLocator`’s ability to predict multiple environmental covariates simultaneously may help retain signal in such cases, though this remains to be tested explicitly. Within the range of architectures tested, `EcoLocator` performed comparably across simulations with clustered and distributed QTL arrangements, and it requires no prior specification of candidate loci or assumptions about the genetic basis of adaptation.

However, several limitations should be noted. First, `EcoLocator` is a supervised method that requires georeferenced samples paired with environmental data for training. Its predictions are therefore bounded by the geographic and environmental range of the training set, and extrapolation to novel environments or unsampled regions should be interpreted cautiously. Second, like all machine learning approaches, the model learns correlations rather than causal mechanisms. High prediction accuracy for an environmental variable does not by itself demonstrate that selection on that variable has shaped genetic variation. As Booker et al. (2024) emphasize, GEA signals do not constitute a test of local adaptation: correlated environmental predictors, spatially structured neutral variation, allelic surfing during historical range expansions, and adaptation to unmeasured but environmentally correlated biotic factors, such as local mycorrhizal or soil microbial communities, can all generate allele frequency patterns that mimic adaptive signals (Klopfstein, Currat, and Excoffier, 2006; Novembre and Di Rienzo, 2009). Isolation by distance alone can inflate genotype–environment correlations when environmental gradients are spatially autocorrelated (Meirmans, 2012). `EcoLocator`’s joint prediction framework does not resolve this fundamental ambiguity; high environmental prediction accuracy could reflect true adaptive signal, demographic history, or both. Third, our simulations use a simplified landscape with a single environmental axis and a modest genome size. Natural systems involve many correlated environmental variables, complex demographic histories, and genomes orders of magnitude larger, all of which may affect model performance in ways not fully captured here.

### Future directions

Several extensions of this work merit exploration. First, our simulations assume a purely additive genetic architecture, and whether prediction accuracy and attribution hold up under non-additive genotype–phenotype maps remains untested.

Second, expanding the environmental complexity of simulations to include multiple correlated environmental axes, more complex spatial covariance among those axes, and temporally varying selection would better approximate the conditions under which `EcoLocator` will be applied in practice. A particularly promising application is using differences in environmental prediction accuracy to identify which environmental gradients are most strongly reflected in genomic variation. Many landscape genomic and genomic-offset approaches consider multiple environmental predictors, but not all environmental axes are expected to contribute equally to local adaptation. Because `EcoLocator` predicts each environmental target separately, variation in prediction accuracy among targets provides information about the strength of genotype–environment correspondence along each axis. For species with limited ecological or common-garden information, this could help researchers and managers prioritize which environmental variables warrant the greatest consideration when designing management strategies, seed-transfer guidelines, or subsequent experimental studies. The same logic extends to geographic prediction: poor geographic accuracy can indicate weak population structure, high gene flow, or limited spatial signal in the sampled genomic data, making prediction accuracy itself a diagnostic signal for species that lack common-garden data or prior characterization.

Third, the framework could also be extended to future climate applications. `EcoLocator`-estimated climates of origin could be compared with projected future conditions to evaluate source–site climate mismatch and identify candidate sources for assisted migration or restoration. This would connect genotype-based inference of historical climate associations with decisions about where biological material may be best suited under changing conditions.

Finally, systematic comparison with a broader suite of GEA methods across a wider range of evolutionary scenarios would help delineate the conditions under which `EcoLocator`’s joint prediction approach offers the greatest advantage over single-task or univariate alternatives.

## Acknowledgments

We thank Peter Ralph and Ángel Rivera-Coĺon for their helpful comments on the manuscript, Amanda de la Torre and Athena Allen (Northern Arizona University) for mapping Axiom SNP markers to a new unpublished Douglas-fir genome (version 5.29), Glenn Howe (Oregon State University) for providing original transcript sequences used in Axiom SNP array design (as well as 500-foot interval seed zone climate summaries for Oregon and Washington), and all members of the Kern-Ralph colab who provided feedback and support throughout this project. This work was supported by NIH grants R35GM148253 and R01HG010774 to ADK and USDA Forest Service Award RDBIL-ERR16 to RCC.

## Supplementary Materials

**Figure S1:**
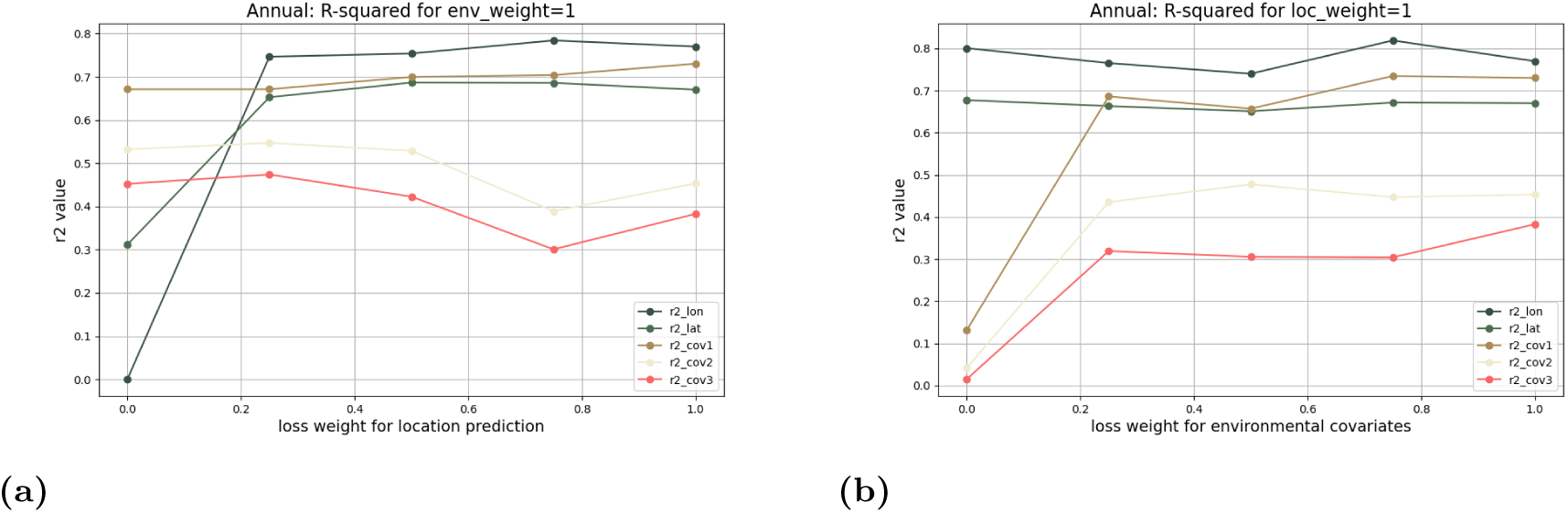
Loss weight cross-validation. To determine the default loss weight value for location and environmental prediction, we tested sliding this value at the command line for each output task (location prediction and environmental covariate prediction) on the empirical Douglas-fir data, while holding the other value at 1. In panel **a)**, we see that when the loss weight is held at a value of 0 for environmental covariate prediction, `EcoLocator` essentially “focuses” entirely on the location prediction task, reflected in the low *R*^2^ scores for all three environmental targets and high prediction scores for location. We see the same effect in the opposite direction in panel **b)**, where we held the loss weight of the environmental output head at 1 and scaled the loss weight for location prediction from 0–1. Again we see that when loss weight for location prediction is held at 0, the model prioritizes predicting on environmental covariates. Reassuringly, this analysis also demonstrates that `EcoLocator` recovers the true values for environmental covariates without over-reliance on the spatial information housed in the location values. Lastly, we see negligible effects on prediction performance in one output or the other when one output head is essentially turned off/down. Because of this, we hold the default value for both prediction tasks at 1.0.

**Figure S2:**
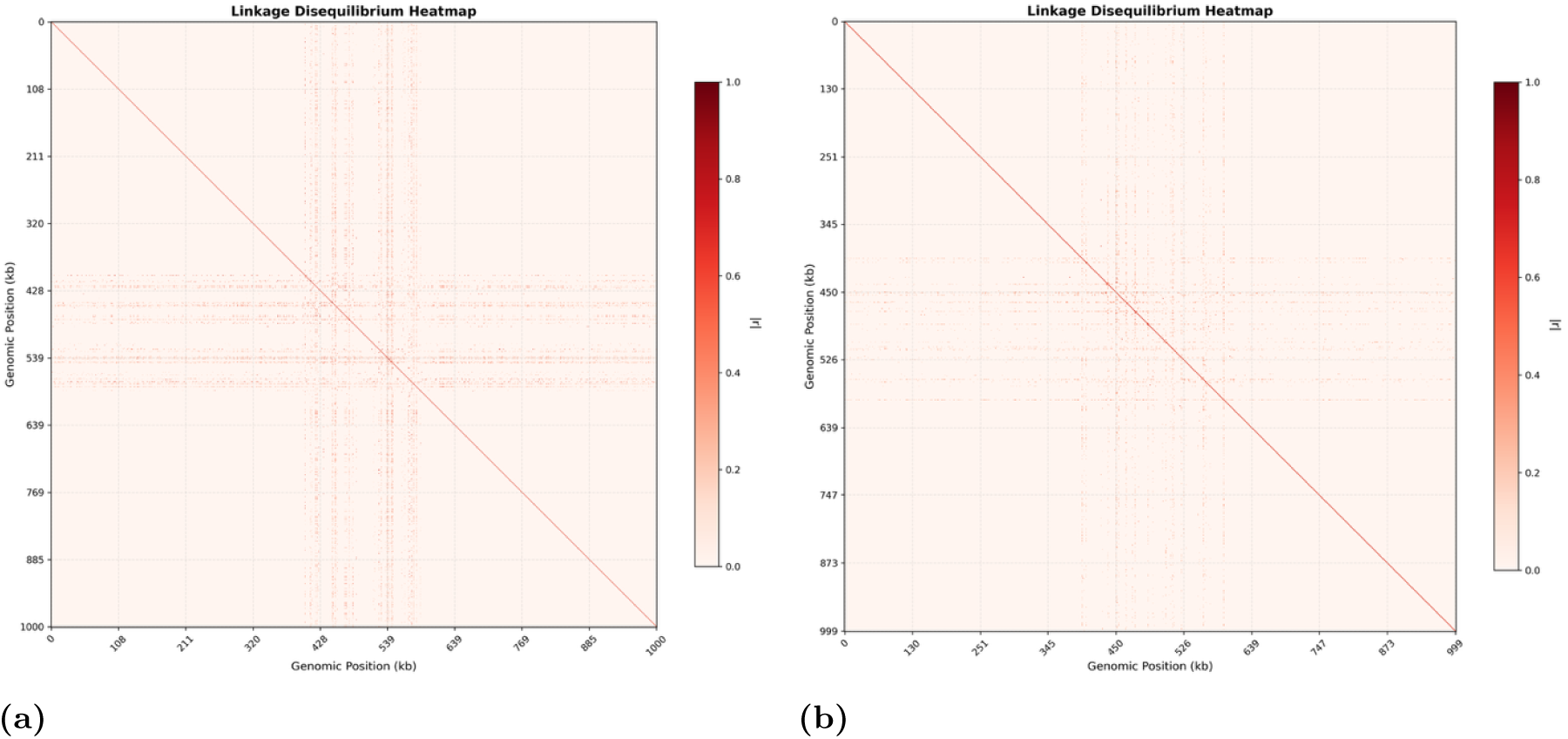
Linkage disequilibrium under different recombination rates for genomic architecture 1. **(a) Recombination** = 10*^−^*^8^; **b) Recombination** = 10*^−^*^6^). Linkage disequilibrium between selected QTL windows and SNPs outside the selected region. To identify SNPs in linkage disequilibrium (LD) with the selected QTL window used in genetic architecture 1, we calculated pairwise correlations between SNPs inside the selected window and SNPs outside the window. These plots do not represent genome-wide LD decay or complete LD structure across the simulated genomic element; rather, they show the subset of outside-window SNPs that pass the LD threshold with SNPs in the selected region. Panel a shows LD partners under lower recombination, where LD extends more broadly away from the selected window. Under this parameterization, 525 SNP pairs showed very strong LD (*r*^2^ *≥* 0.8), 1,026 showed strong LD (*r*^2^ *≥* 0.5), and 2,923 showed moderate LD (*r*^2^ *≥* 0.2). Panel **b** shows the same analysis under higher recombination, where LD with the selected window is reduced. Under this parameterization, 9 SNP pairs showed very strong LD (*r*^2^ *≥* 0.8), 24 showed strong LD (*r*^2^ *≥* 0.5), and 309 showed moderate LD (*r*^2^ *≥* 0.2). Because SNPs were filtered to include only LD partners of the selected QTL window, long-range LD in these panels should be interpreted as evidence of SNPs that remain correlated with the selected region, rather than as a general description of background LD across the genome.

**Figure S3:**
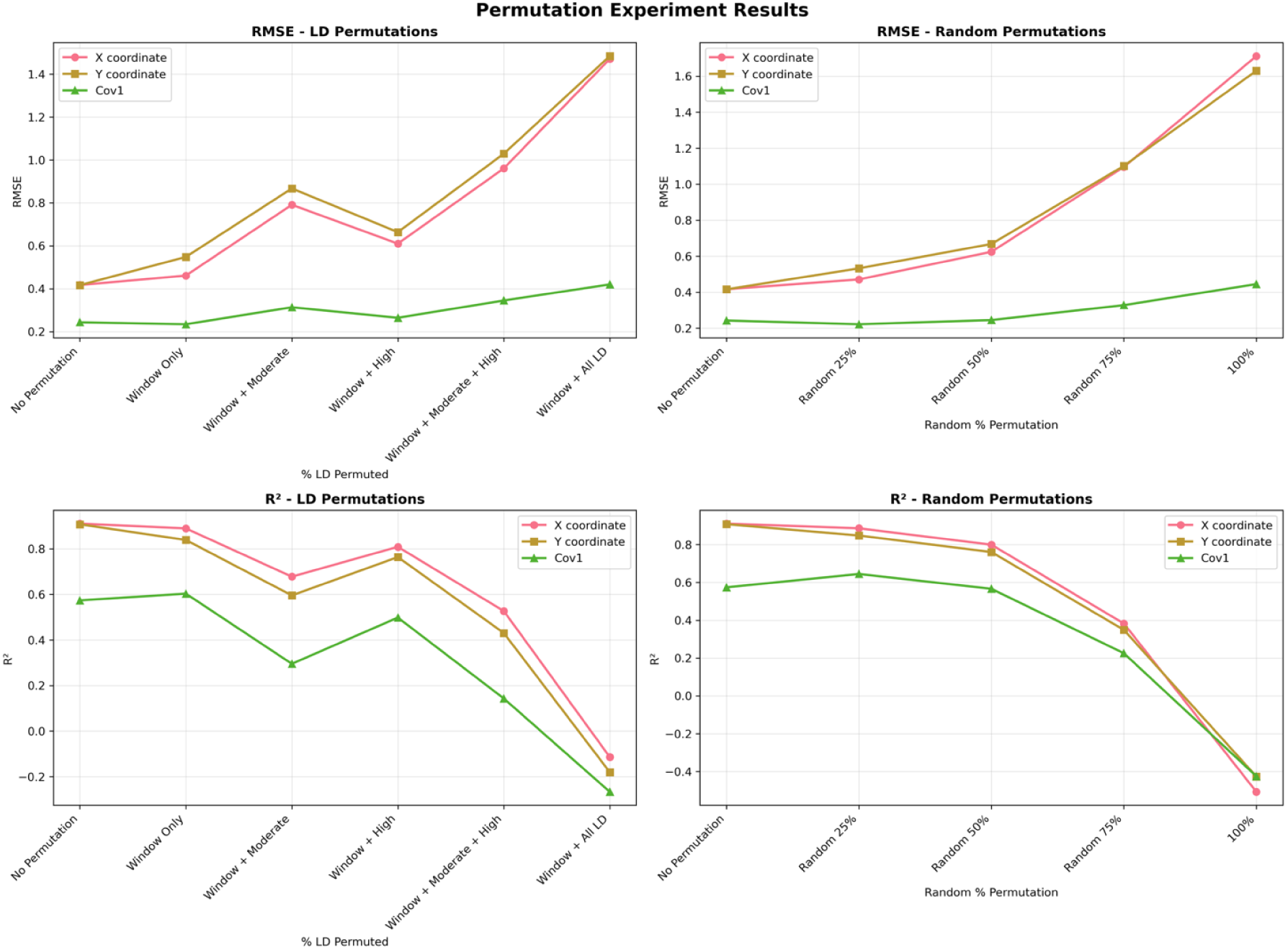
RMSE and *R*^2^ of EcoLocator predictions under different permutation regimes (combinations of random % permuted and sites in LD permuted). `EcoLocator` prediction performance under random and LD-targeted permutation regimes (*R*^2^ is computed out-of-sample and can be negative when predictions are worse than the training-set mean). To assess whether prediction accuracy depends primarily on the selected QTL window, SNPs in linkage disequilibrium with that window, or genome-wide signal distributed across the genotype matrix, we permuted subsets of the genotype matrix across individuals prior to prediction. The left column shows targeted permutations of the selected window alone and the selected window combined with SNPs in moderate, high, or all LD with the selected region. The right column shows random genome-wide permutations of increasing proportions of SNPs. The top row shows root mean squared error (RMSE), and the bottom row shows *R*^2^ for X coordinate (longitude), Y coordinate (latitude), and the environmental covariate. In both permutation designs, prediction accuracy declines as more genotype information is shuffled, indicating that `EcoLocator` relies on genetic signal distributed across the genotype matrix. However, permuting the selected window together with all LD-linked sites causes a sharp loss of performance, suggesting that SNPs correlated with the selected region carry important predictive information. Randomly permuting all SNPs produces similarly poor performance, confirming that prediction accuracy is lost when genotype–environment and genotype–location associations are fully disrupted. Note that the “window + all LD” grouping comprised roughly 75% of sites, so comparing that to the random 75% permutation again reiterates the importance of SNPs in LD.

**Figure S4:**
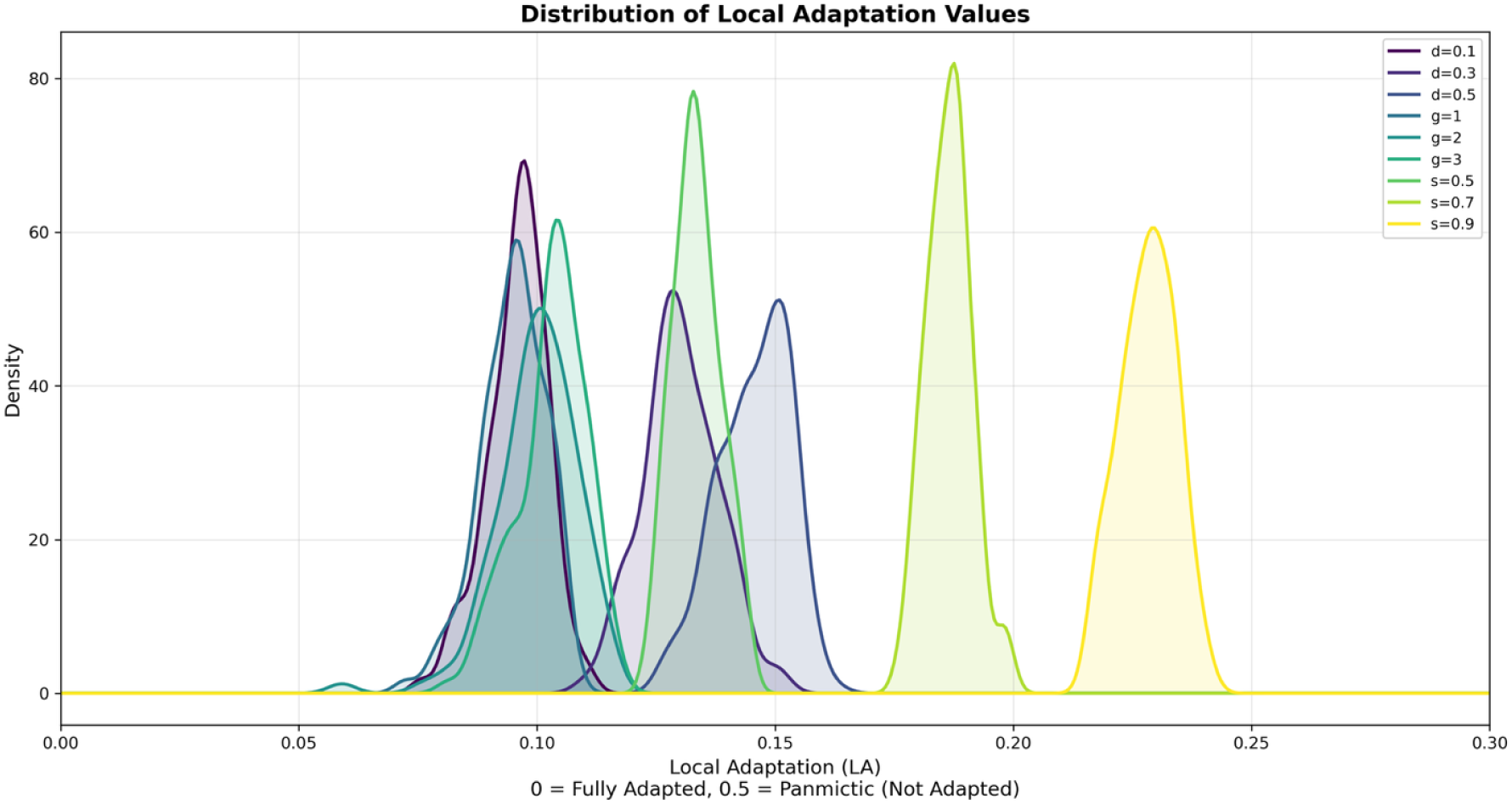
Distribution of strength of local adaptation across parameter space. Kernel density estimates of local adaptation (LA) values for 100 simulation replicates per parameter combination across three parameter explorations on an 8×8 grid-like landscape with Hurst = 0 (no autocorrelation between space and environment). LA is calculated as the mean absolute difference between individual phenotypic values and their local optima; lower values indicate stronger local adaptation. The selection parameter *σ_K_* controls the standard deviation of the fitness function, where smaller values produce a tighter fitness distribution and thus stronger selection. Dispersal (*σ_D_* = 0.1, 0.3, 0.5; genetic architecture 1 and *σ_K_*= 0.5 fixed): increasing dispersal progressively eroded local adaptation, with mean LA values of 0.096, 0.130, and 0.146 for *σ_D_* = 0.1, 0.3, and 0.5, respectively. Genetic architecture (*g* = 1, 2, 3; *σ_D_* = 0.1 and *σ_K_* = 0.5 fixed): distributing QTLs increasingly across the genomic element had a modest effect on local adaptation, with mean LA values of 0.095, 0.100, and 0.103 for *g* = 1, 2, and 3, respectively. Selection (*σ_K_* = 0.5, 0.7, 0.9; genetic architecture 1 and *σ_D_*= 0.3 fixed): an extended range of selection strengths was used here to illustrate how local adaptation varies across increasingly weak selection regimes. Weaker selection substantially reduced local adaptation, with mean LA values of 0.134, 0.187, and 0.228 for *σ_K_*= 0.5, 0.7, and 0.9, respectively.

**Figure S5:**
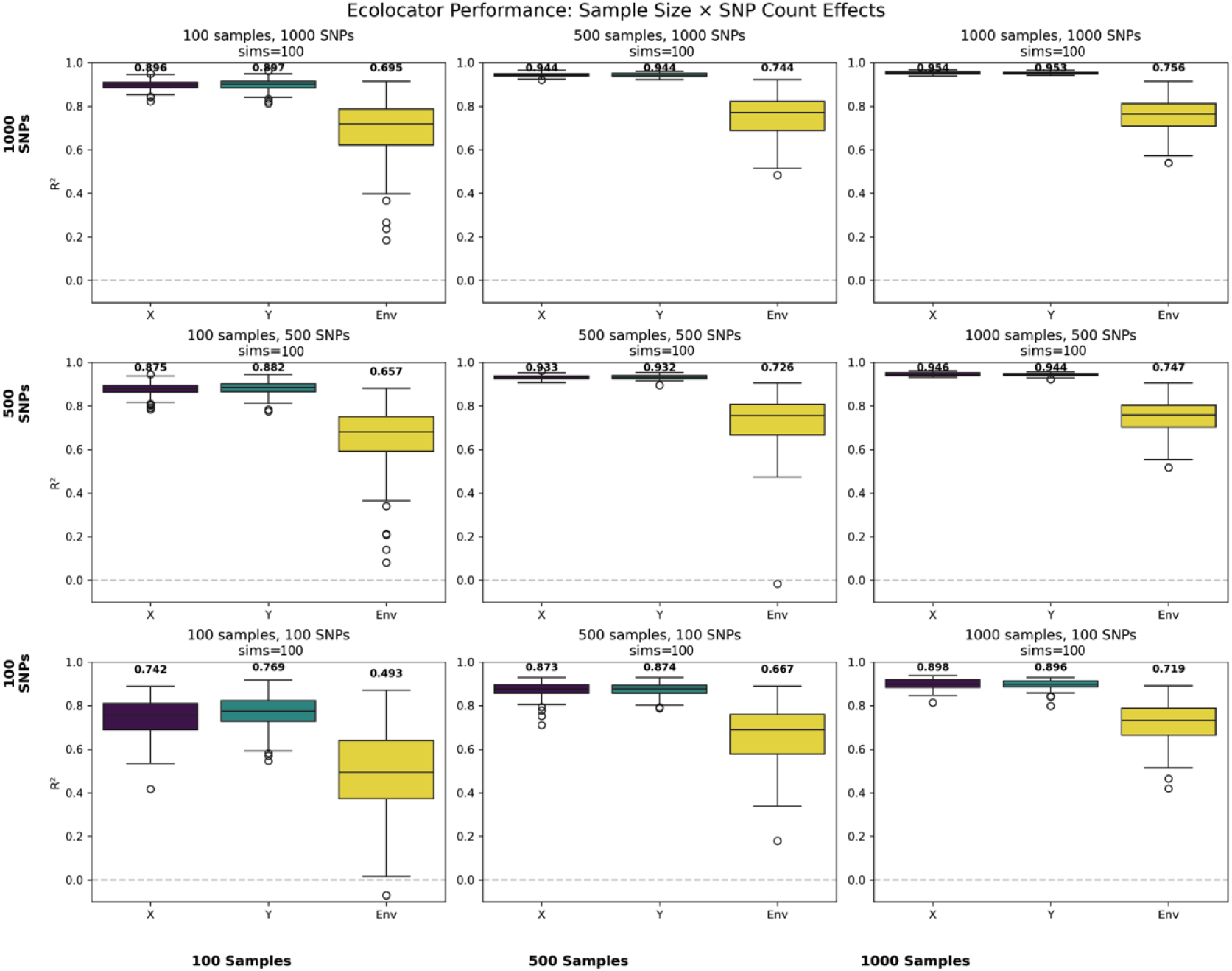
Effects of down-sampling individuals and SNPs as input to EcoLocator. `EcoLocator` performance under various downsampling regimes of the same simulated data (*σ_D_* = 0.1, *σ_K_* = 0.5, *g* = 1). In the top row, there are 1000 SNPs made available to the model; in the middle and bottom rows, there are 500 and 100 SNPs respectively. In the columns we vary sample size, with 100, 500, and 1000 samples from left to right. *R*^2^ values across 100 simulation replicates are summarized in boxplots, with Longitude (x) in purple, Latitude (y) in teal, and Environment in yellow. Because `EcoLocator` learns from variation present in the data, it is no surprise that we observe the highest prediction accuracy across targets (Lon. *R*^2^ = 0.954, Lat. *R*^2^ = 0.953, and Env. *R*^2^ = 0.756) with 1000 samples and 1000 SNPs (top right corner), and we see the lowest prediction accuracy (Lon. *R*^2^ = 0.742, Lat. *R*^2^ = 0.769, and Env. *R*^2^ = 0.493) with only 100 SNPs and 100 samples (bottom left corner). Under the sample sizes and SNP sets explored here, we do not observe diminishing returns with “too much” data being shown to the model, and similarly, we do not see a complete absence of predictive performance when sample size and SNP set are restricted.

**Figure S6:**
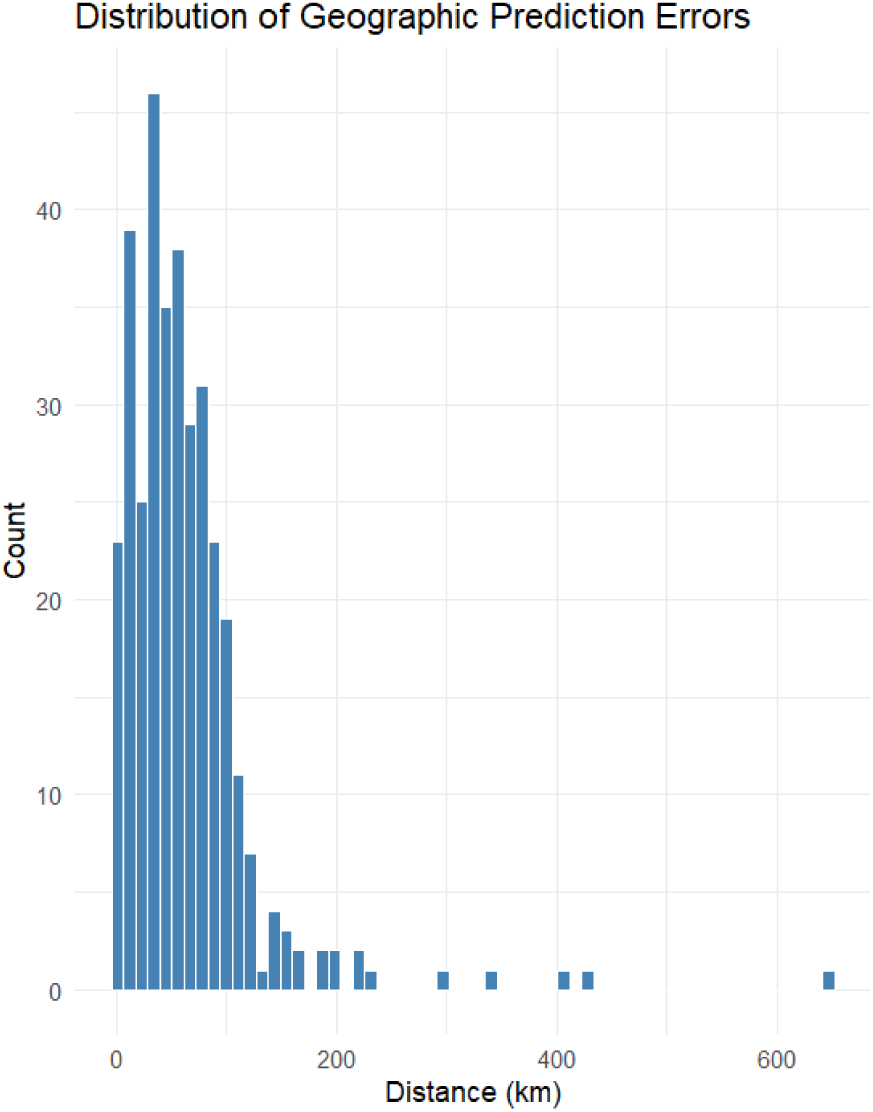
Distribution of geographic prediction errors on Douglas-fir data. A histogram of geographic distance prediction errors in our empirical application of `EcoLocator` on the Douglas-fir case study data. Distance in kilometers (km) is on the x-axis and count is on the y-axis. Most errors fall between 0-100km, with others as high as *>*600km.

**Figure S7:**
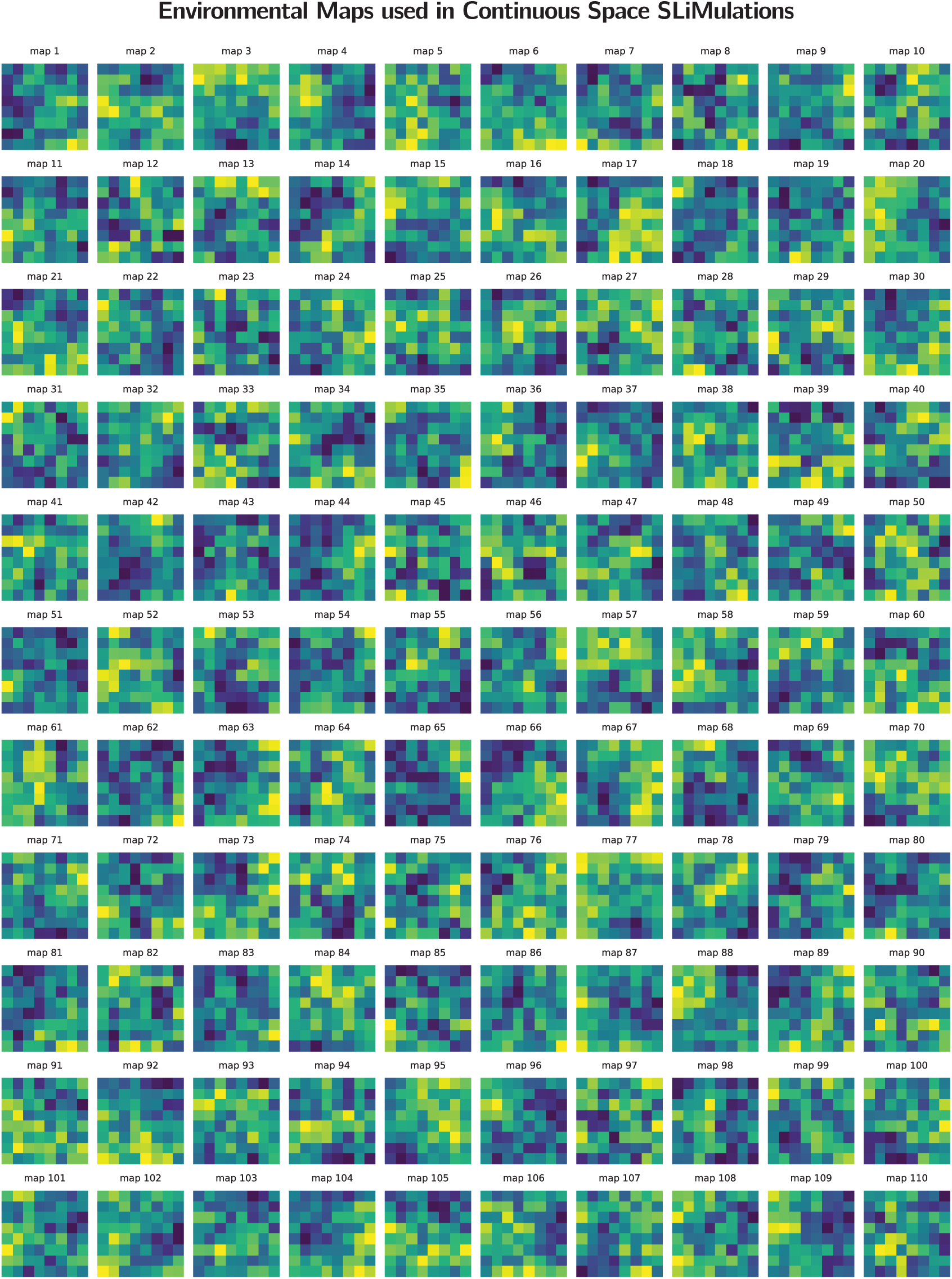
Environmental Maps used in Continuous Space. SLiM **simulations.** Environmental maps generated with the NLMpy python package with pixel count set to 8×8 and Hurst (autocorrelation) set to H=0. We generated 110 maps and sampled 100 of them randomly for each simulation regime.

**Figure S8:**
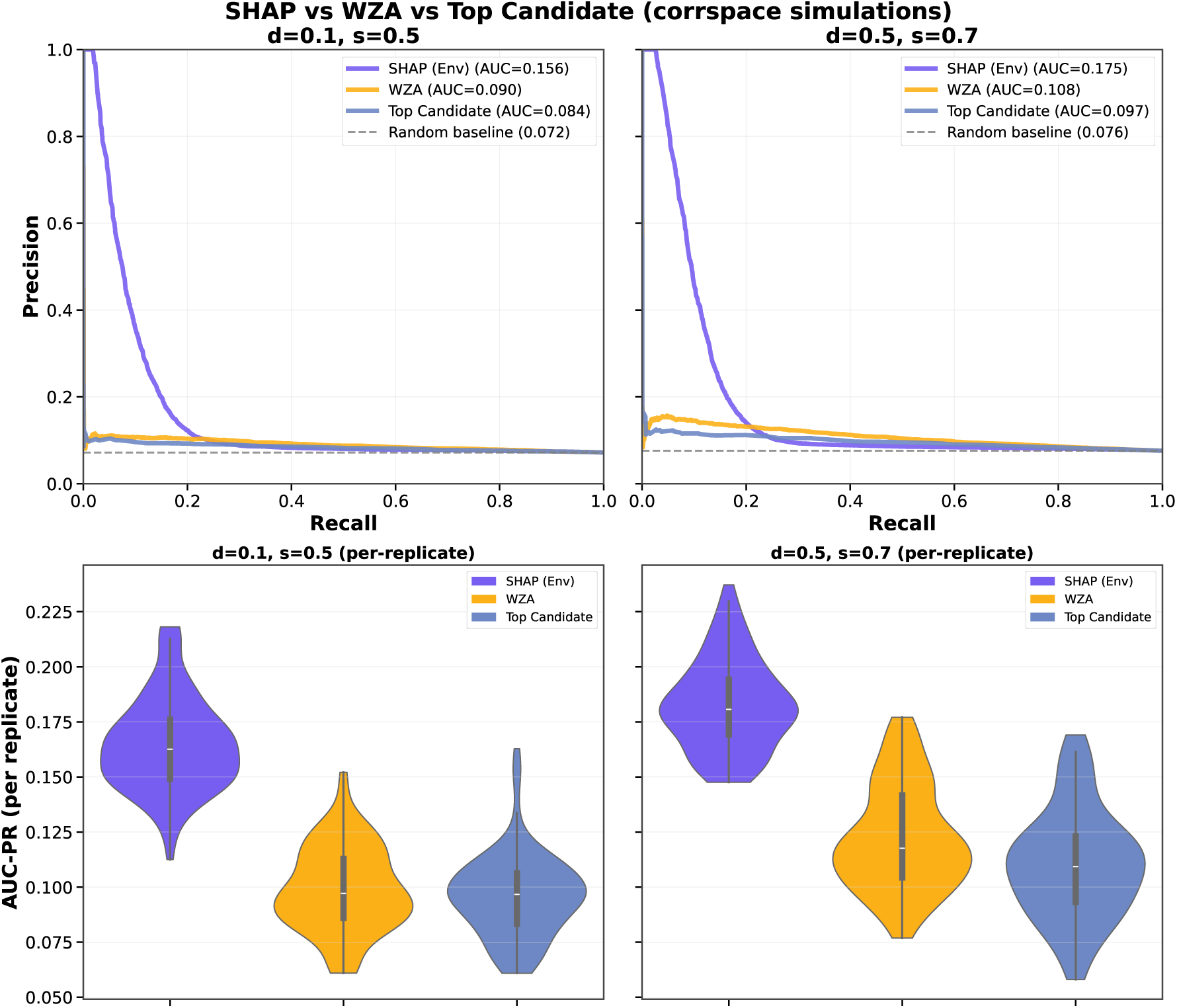
AUC-PR of EcoLocator ×SHAP, WZA, and Top-Candidate GEA Methods Across Two Dispersal/Selection Regimes. Precision-recall curves (top) and per-replicate AUC-PR violin plots (bottom) comparing three genotype-environment association methods for identifying adaptive SNPs: `EcoLocator` ×SHAP (purple; mean *|*SHAP*|* for the environmental covariate, ranked by an empirical FDR score computed from known ground-truth adaptive status), `WZA` (yellow), and top-candidate (blue). Top row: precision-recall curves pooling all SNPs across 100 simulation replicates per regime, scored by *−* log_10_ of each method’s *p*-value or empirical FDR score against true adaptive status; dashed line = random-baseline AUC-PR (proportion of adaptive SNPs). Bottom row: distribution of AUC-PR computed separately within each of the 100 replicates, showing the consistency of each method’s performance across independent simulations. Left: limited-dispersal, strong-selection regime (dispersal=0.1, std. dev. of fitness function=0.5). Right: higher-dispersal, weaker-selection regime (dispersal=0.5, std. dev. of fitness function=0.7). `EcoLocator` ×SHAP outperforms `WZA` and top-candidate under both regimes, both pooled and at the per-replicate level.

**Figure S9:**
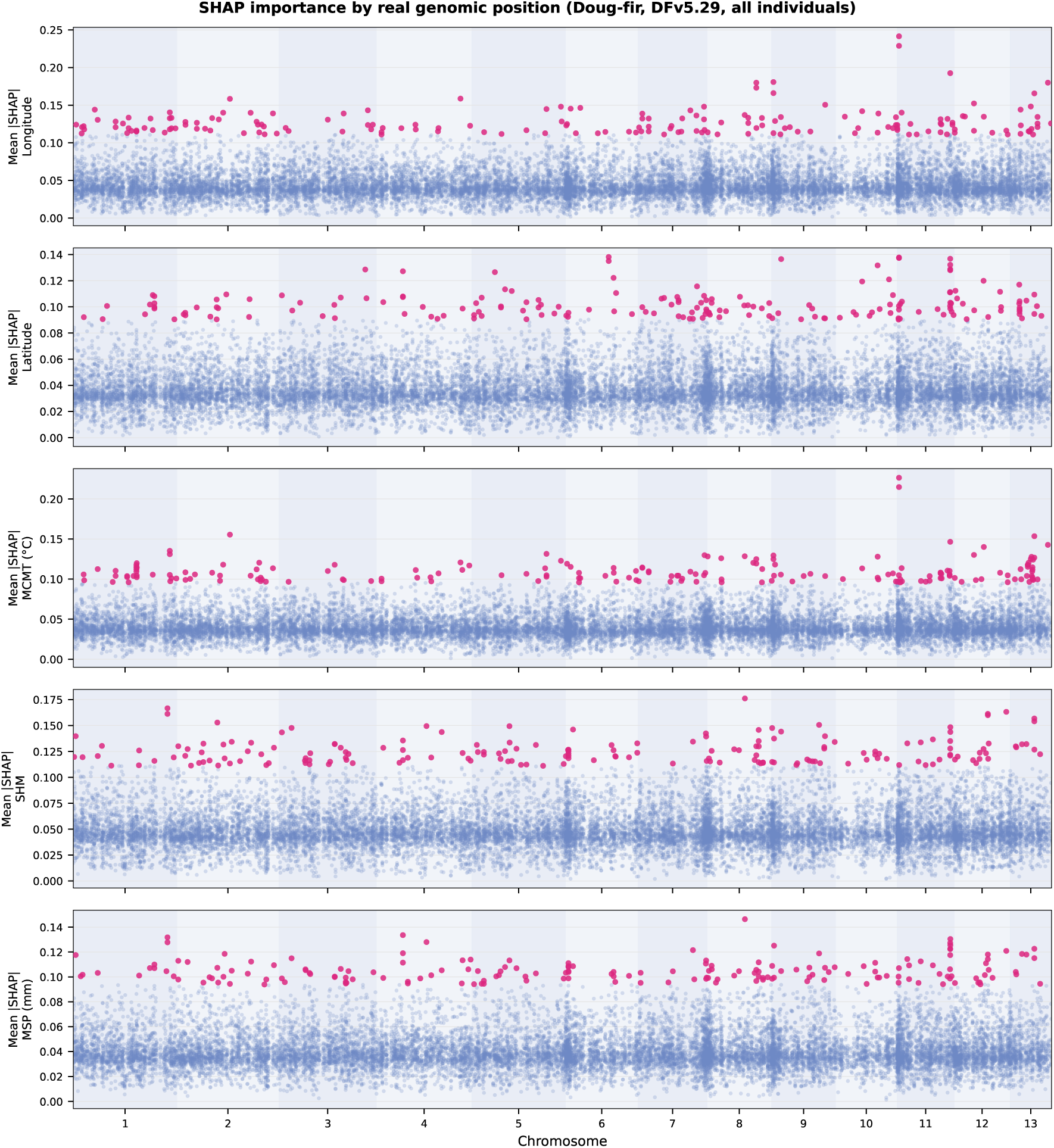
Mean *|*SHAP*|* by Real Genomic Position Across Five Prediction Targets for Douglas-fir. Manhattan-style plots of mean *|*SHAP*|* (averaged across leave-one-out cross-validation folds) at real chromosomal positions (chromosomes 1–13, Douglas-fir reference assembly DFv5.29; SNP positions resolved by SAM alignment restricted to primary, high-confidence mappings, MAPQ *≥* 30) for five prediction targets, top to bottom: longitude, latitude, Mean Cold Month Temperature (MCMT), Summer Heat:Moisture Index (SHM), and mean summer precipitation (MSP). SNPs in the 179 highest-attribution SNPs (approximately the top 1%) of mean *|*SHAP*|* for a given target are highlighted in pink; all other SNPs are shown in blue. Alternating background shading demarcates chromosome boundaries.

**Figure S10:**
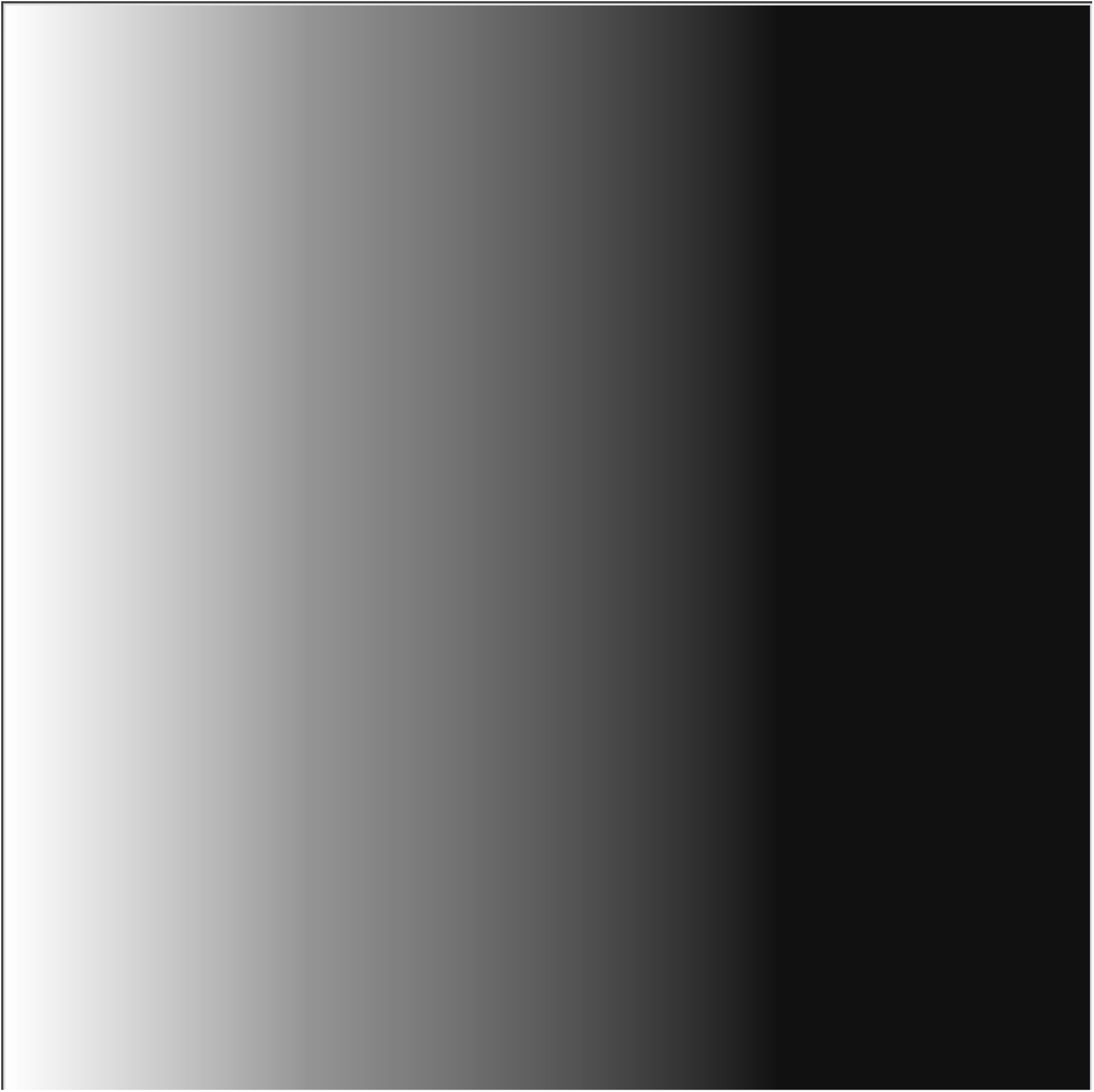
Environmental Map used in Correlated Space Simulations. A grayscale left-right gradient image used in simulations to represent an arbitrary environmental variable correlated with space along the longitudinal axis.

**Figure S11:**
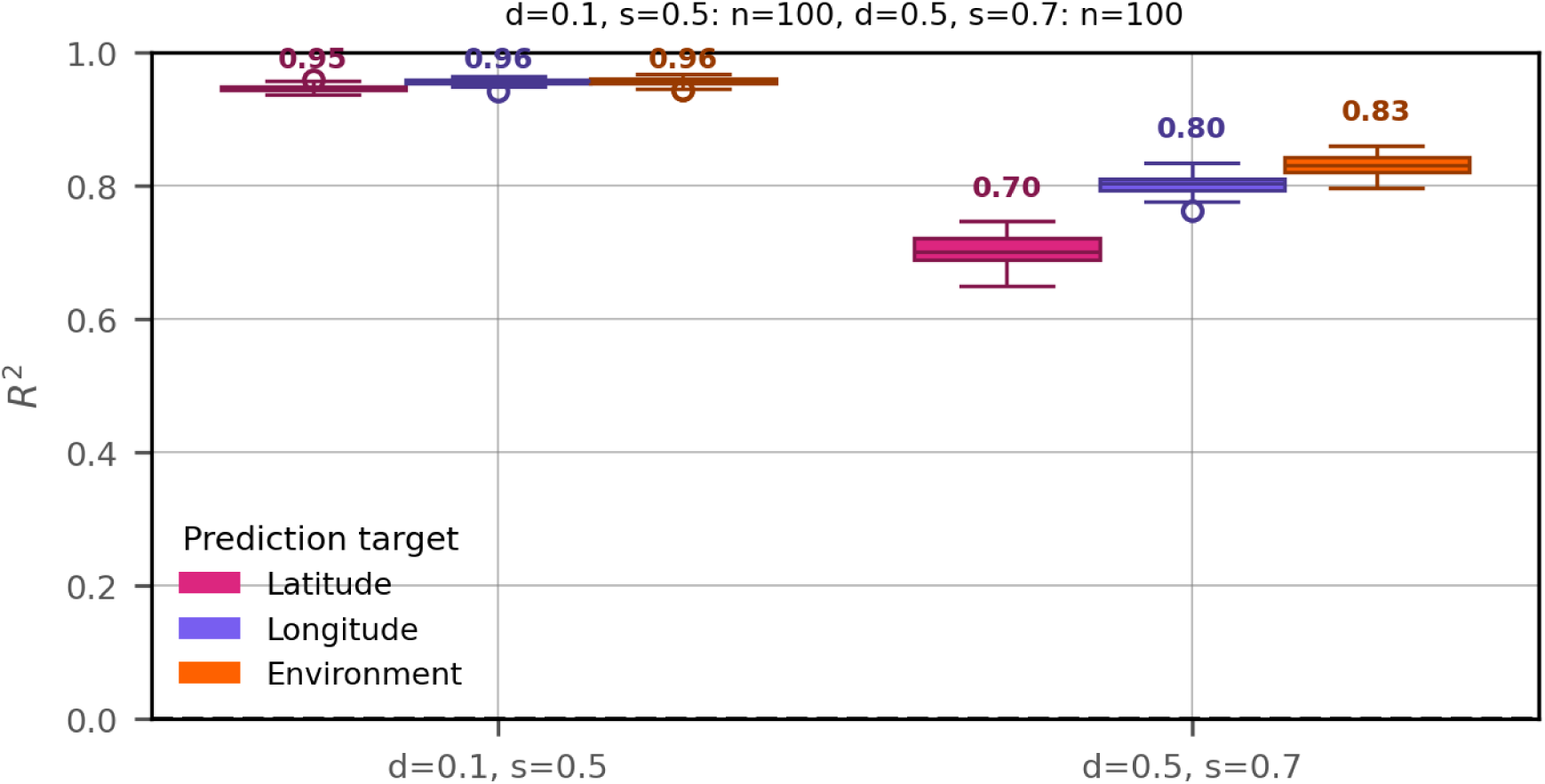
Per-replicate LOOCV prediction accuracy (*R*^2^) across two dispersal/selection regimes. For each of 100 independent simulation replicates per regime, predictions from all leave-one-out cross-validation folds were pooled and compared to the known sampled values to compute a single *R*^2^ per replicate for each of three prediction targets: latitude, longitude, and environment (cov1). Boxplots show the distribution of these per-replicate *R*^2^ values (box = interquartile range, line = median, whiskers = 1.5×IQR, points beyond whiskers = outlier replicates); numbers above each box give the across-replicate mean *R*^2^. Left: limited-dispersal, strong-selection regime (*σ_D_*=0.1, *σ_K_*=0.5). Right: higher-dispersal, weaker-selection regime (*σ_D_*=0.5, *σ_K_*=0.7). Prediction accuracy is uniformly high and tightly distributed for all three targets under limited dispersal/strong selection (mean *R*^2^=0.95–0.96), but declines and becomes more variable across replicates under higher dispersal/weaker selection, most notably for latitude (mean *R*^2^=0.70 vs. 0.80–0.83 for longitude and environment).

**Figure S12:**
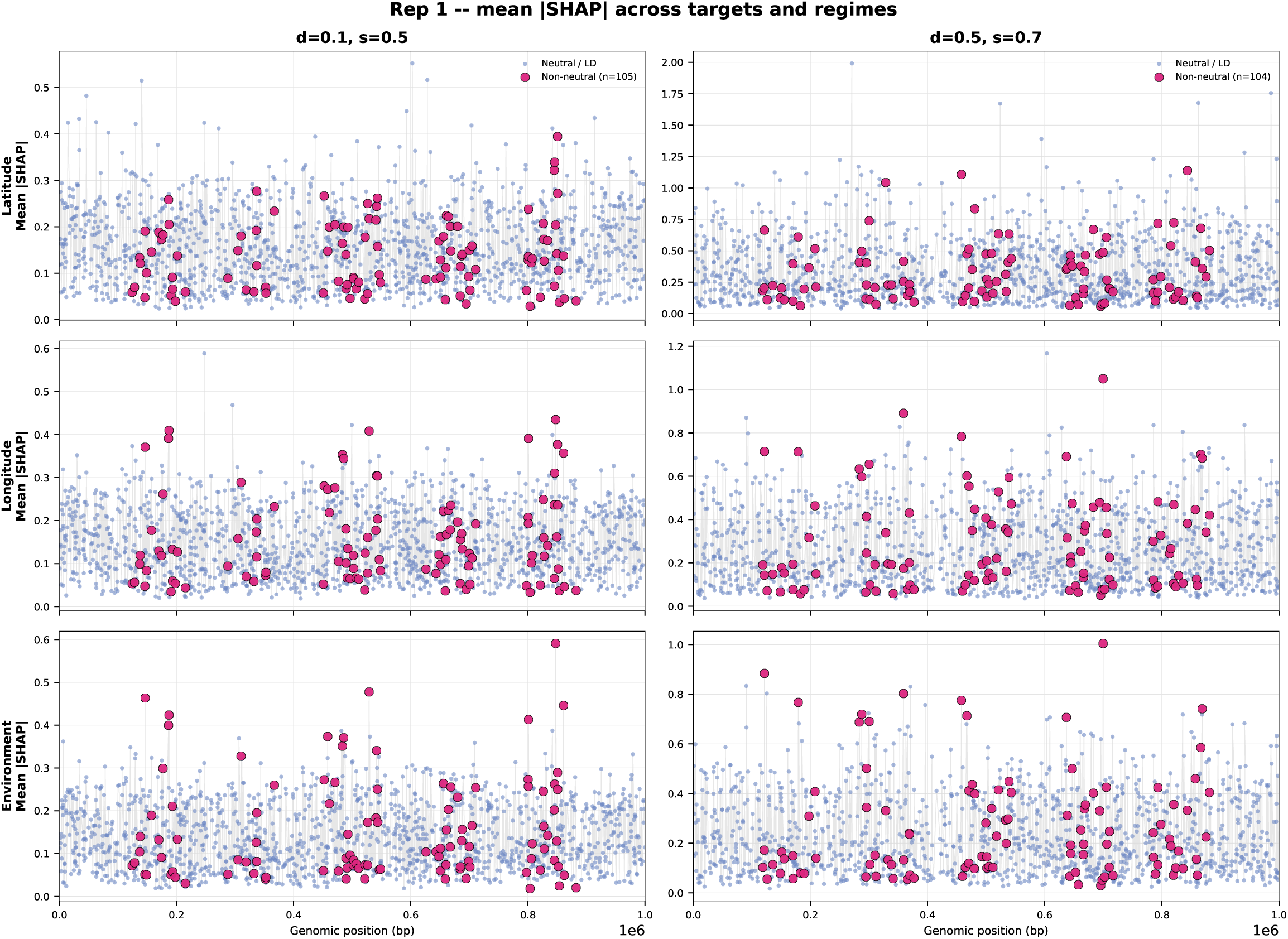
Genomic Distribution of Mean *|*SHAP*|* Across Three Prediction Targets in a Representative Replicate. Mean *|*SHAP*|* by genomic position (bp) for one representative simulation replicate (rep1), shown for three prediction targets (rows: latitude, longitude, environment) and both dispersal/selection regimes (columns: limited-dispersal, strong-selection, dispersal=0.1, std. dev. of fitness function=0.5, left; higher-dispersal, weaker-selection, dispersal=0.5, std. dev. of fitness function=0.7, right). SNPs classified as non-neutral (true causal loci; magenta, black-outlined) are overlaid on all other neutral/LD-linked SNPs (light blue).

**Table S1:** *R*^2^ values for EcoLocator predictions versus Location+ClimateNA estimates. For latitude and longitude, values are `EcoLocator`’s leave-one-out *R*^2^, duplicated here for reference (ClimateNA does not predict location, hence “–”). For the three climate variables, *R*^2^ is shown for `EcoLocator`’s direct climate prediction and for ClimateNA modeled values at `EcoLocator`’s predicted locations, computed for the same 348 individuals from a single leave-one-out run; *p*-values are from a paired Wilcoxon signed-rank test on absolute error (direct vs. Location+ClimateNA).

| Predicted Location and Climate | Geolocation and Climate Prediction | Geolocation and ClimateNA | $p$-value |
| --- | --- | --- | --- |
| Latitude (LAT, °) | 0.83 | – | – |
| Longitude (LON, °) | 0.75 | – | – |
| Mean Cold Month Temperature (MCMT, °C) | 0.72 | 0.56 | $1.1 \times 10^{-6}$ |
| Mean Summer Precipitation (MSP, mm) | 0.52 | –0.005 | $6.0 \times 10^{-10}$ |
| Summer Heat:Moisture Index (SHM) | 0.54 | 0.005 | $1.6 \times 10^{-13}$ |

**Table S2:**
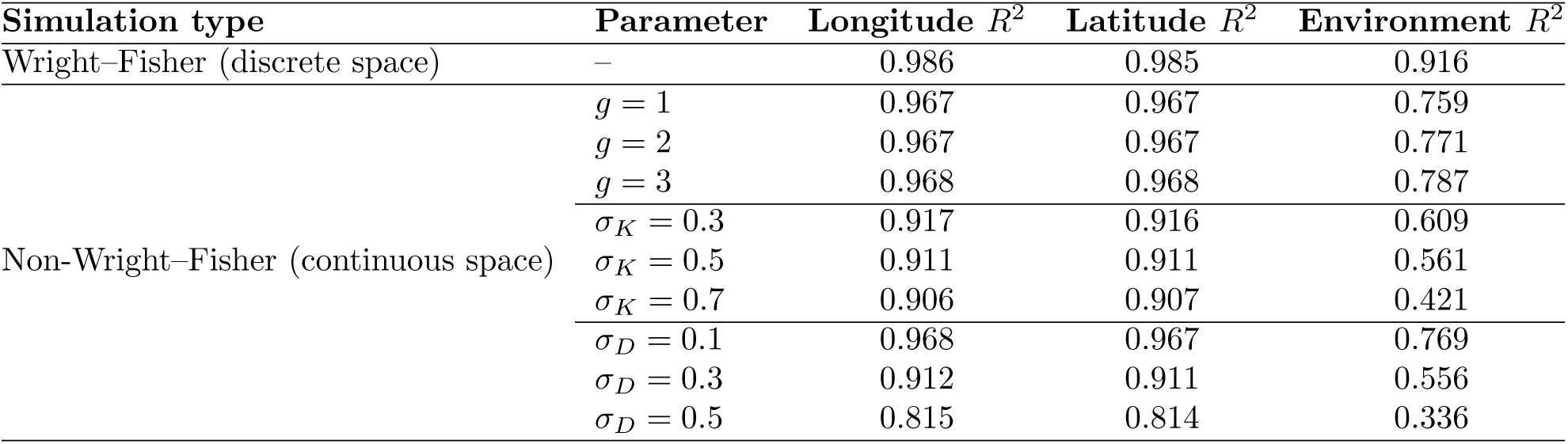
Mean *R*^2^ values for EcoLocator predictions of longitude, latitude, and environment across simulation types.

**Table S3:** Pairwise overlap among high-attribution SNPs across EcoLocator prediction targets in coastal Douglas-fir.

| Prediction target | LAT | LON | MCMT | MSP | SHM |
| --- | --- | --- | --- | --- | --- |
| Latitude (LAT) | 179 | 10 (5.6%) | 13 (7.3%) | 33 (18.4%) | 31 (17.3%) |
| Longitude (LON) |  | 179 | 97 (54.2%) | 5 (2.8%) | 4 (2.2%) |
| MCMT |  |  | 179 | 5 (2.8%) | 5 (2.8%) |
| MSP |  |  |  | 179 | 133 (74.3%) |
| SHM |  |  |  |  | 179 |

## Footnotes

1 Source code, installation instructions, and vignettes are available at https://github.com/kr-colab/ecolocator and https://ecolocator.readthedocs.io/en/latest/.

